# A deterministic computational kernel encoded in the human genome

**DOI:** 10.64898/2026.04.12.718009

**Authors:** Jasmine Levy

## Abstract

A computational kernel is defined by four properties: initialization from raw input, a fixed instruction set, a process table with memory organization, and inter-component signal dispatch. Here we show that the human genome satisfies all four. Applying a deterministic 6-bit encoding to the complete human proteome (83,587 isoforms from 32,281 genes) and all chromosome assemblies, we extract 1,932 recurring byte-level vocabulary patterns mapping to 27 functional categories, identify 4,936 genome programs with chromosome-based memory segments, and trace a dispatch network of 543,554 edges routed through a single relay hub. All seven network entry points converge on the mitochondrial genome, which functions as a read-only boot origin. Five null model tests, fifteen robustness analyses, and independent validation against DepMap gene essentiality and STRING protein-protein interactions confirm that this architecture is a property of the genome itself. Vocabulary conservation tracks evolutionary divergence across nine species. The same pipeline applied to composition-matched random sequences fails all four properties.

## Introduction

In computer science, a kernel is the irreducible core of an operating system, defined by four properties: it boots from raw data without external configuration, it manages a fixed instruction set, it maintains a process table with memory protection, and it dispatches signals between components^22^. These properties are substrate-independent: a system implemented in silicon, vacuum tubes, or biological molecules is a kernel if and only if it satisfies all four criteria. Genomes are routinely described using the language of information: genes are “read,” sequences are “transcribed,” and regulatory elements “control” expression. Recent work has advanced our ability to predict genomic outputs from sequence^1,2,3^, and studies of chromatin architecture have revealed computational properties in genome geometry^4^. Network-level comparisons have drawn explicit parallels between genome regulation and operating system architecture^5^. DNA itself has been shown to function as a digital storage medium using simplified binary encodings^19^, and automated design tools have demonstrated that engineered Boolean logic circuits can be compiled into DNA sequences that execute specified computations in living cells^21^. Yet the computational metaphor applied to the native linear sequence itself has remained just that: no study has demonstrated that a genome’s nucleotide sequence contains the structural components of a computational system, including an instruction set, a process table, a dispatch network, and a boot sequence that initializes from raw data. Here we present evidence that a deterministic encoding of the human genome reveals such architecture.

We apply a 6-bit encoding pipeline to the complete human proteome (83,587 protein isoforms from 32,281 genes) and full chromosome assemblies^6^, extracting a computational vocabulary of 1,932 recurring byte-level patterns. The pipeline encodes each amino acid as a 6-bit value (equivalently, each codon’s three nucleotides contribute 3 × 2 = 6 bits), concatenates these into a bitstream, and reads back 8-bit bytes; the vocabulary patterns therefore operate at byte granularity even though the underlying encoding uses 6-bit residue codes. When encoding proteins (as opposed to raw genomic DNA), each amino acid is deterministically backtranslated to a single representative codon; the resulting byte streams therefore approximate, but do not exactly reproduce, the native genomic coding sequences. These patterns serve as an instruction set: each word maps deterministically to one of 27 functional departments through enrichment analysis against Gene Ontology annotations. Applying the same encoding to chromosomal DNA identifies contiguous regions of elevated vocabulary density, referred to as genome programs, from which recurring primitives and an inter-chromosomal dispatch network emerge. The complete system is assembled as a kernel (Fig. 1) that boots from raw data files, discovers chromosomal roles, builds a process table, and dispatches signals from entry points on the mitochondrial genome through a discovered relay hub (chromosome 19) to target processes genome-wide.

**Figure 1.**
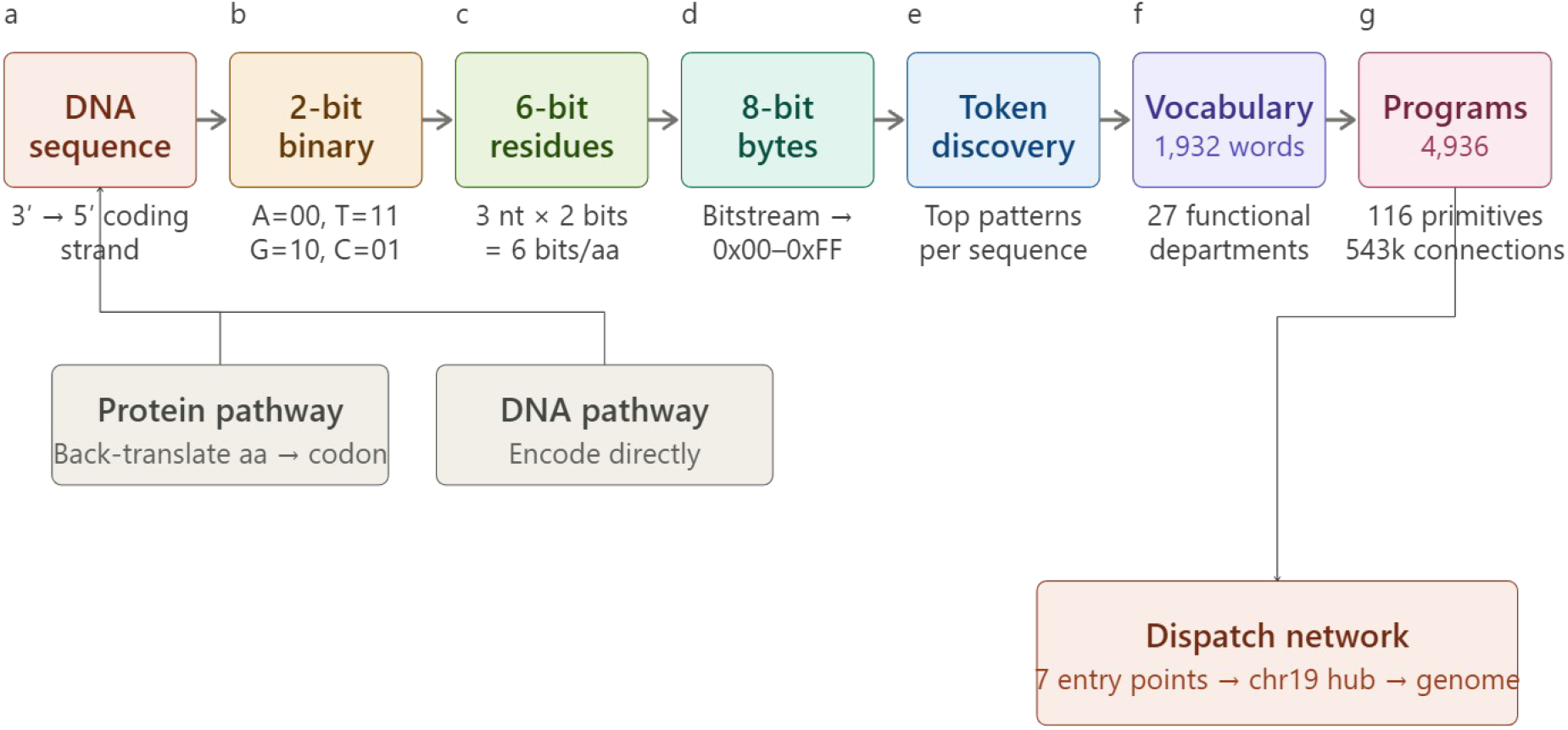
Encoding pipeline overview. DNA coding sequences are converted to byte streams through seven deterministic stages: nucleotide-to-binary encoding (Step 1), binary-to-byte conversion (Step 2), 6-bit residue framing (Step 3), 8-bit byte extraction (Step 4), token discovery (Step 5), functional enrichment into 27 vocabulary departments (Step 6), and program/dispatch assembly (Step 7). The same pipeline applies to both protein sequences (via deterministic back-translation) and raw chromosomal DNA.

The claim that a system satisfies a formal definition requires satisfaction proof: each defining property must be demonstrated and validated against appropriate null models. We address this through a formal boot sequence modelled on hardware power-on self-test (POST) conventions and five null model validation tests assessing encoding specificity, functional convergence, predictive generalization, program recurrence, dispatch hub structure, and cross-species conservation.

This paper is restricted to validating the kernel itself. Applications that build upon the kernel will be addressed in subsequent work.

## Results

### Property 1: The genome boots from raw data

The kernel was booted from the human reference genome using the deterministic encoding pipeline described in Methods (Steps 1-8). The boot sequence completed all five POST phases without failure: 25 chromosomes were enumerated, 116 primitives loaded as the instruction set, 4,936 programs loaded as process images, and 7 entry points identified. Entry points are defined as vocabulary patterns that initiate signal dispatch cascades; in the boot sequence, these are identified at program boundaries on the mitochondrial genome, though the concept generalizes to any chromosome serving as a signal origin (see Methods, Step 7). All seven entry points converge on chrM, a data-driven result, not a design choice. chrM hosts zero entries in the nuclear process table. Circular scanning of the mitochondrial genome, which requires separate treatment due to its closed-circular topology, identifies 136 vocabulary-containing windows (127 recurring, 9 singleton), with 84.6% of the 4,142 scanned positions containing vocabulary from the protein-derived dictionary. This confirms that the computational vocabulary extends to the mitochondrial genome, though chrM’s encoding structure differs from nuclear chromosomes: it lacks the repetitive patterns detectable by frequency analysis and instead encodes information through position-specific vocabulary placement., and gap analysis of cross-chromosome edge ratios independently classifies it as read-only. Its dispatch traffic is balanced and modest (1,135 edges in each direction), consistent with an origin rather than a hub. Gap analysis classified chr19 as the sole relay hub (outbound/inbound ratio 1.55, p = 0.027) and chr9, chrX, and chrY as terminal effector chromosomes, which are chromosomes that receive dispatch signals but do not relay them to other chromosomes (Table 1). The remaining 20 nuclear chromosomes were classified as dual-role (relay-effector). The process table contained 3,069 active processes distributed across 25 memory segments with role-based protection levels.

**Table 1.**
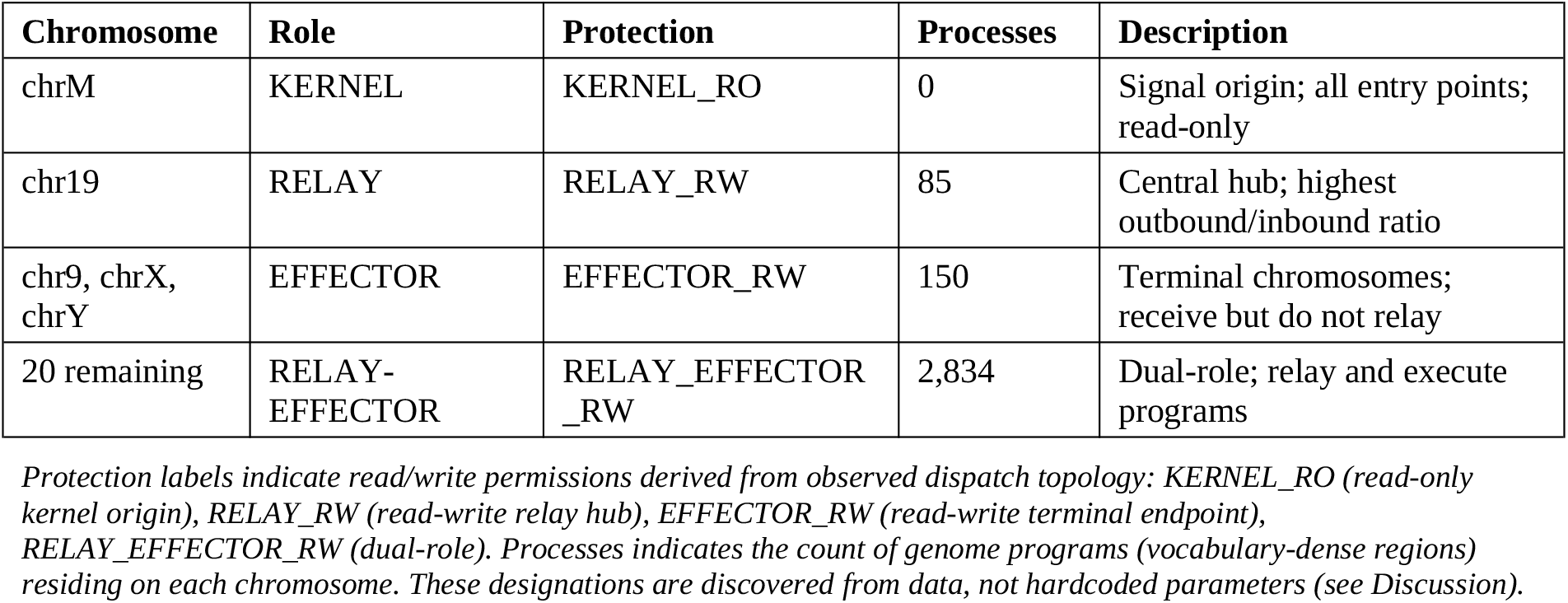
Chromosome role classification discovered by POST.

| Chromosome | Role | Protection | Processes | Description |
| --- | --- | --- | --- | --- |
| chrM | KERNEL | KERNEL_RO | 0 | Signal origin; all entry points; read-only |
| chr19 | RELAY | RELAY_RW | 85 | Central hub; highest outbound/inbound ratio |
| chr9, chrX, chrY | EFFECTOR | EFFECTOR_RW | 150 | Terminal chromosomes; receive but do not relay |
| 20 remaining | RELAY-EFFECTOR | RELAY_EFFECTOR_RW | 2,834 | Dual-role; relay and execute programs |

*Protection labels indicate read/write permissions derived from observed dispatch topology: KERNEL\_RO (read-only kernel origin), RELAY\_RW (read-write relay hub), EFFECTOR\_RW (read-write terminal endpoint), RELAY\_EFFECTOR\_RW (dual-role). Processes indicates the count of genome programs (vocabulary-dense regions) residing on each chromosome. These designations are discovered from data, not hardcoded parameters (see Discussion).*

Signal dispatch traced 543,554 edges from the 7 mitochondrial entry points through the network, with chrM:0x575D generating 93% of all signals routed via the chr19 relay hub. These 543,554 edges represent individual vocabulary-pattern co-occurrences across chromosomes (hop-1 direct matches); aggregating by source and target chromosome yields the 552 directed inter-chromosome connections. The full dispatch graph, defined as the mathematical structure of edges and nodes (552 inter-chromosome connections), extends to hop-2 (12.17 million edges), provided in the repository but not analyzed here. No parameters were configured manually; chromosome roles, relay hubs, and effector assignments were discovered entirely from data.

### Property 2: The encoding reveals a structured instruction set

Application of the 6-bit encoding pipeline to the complete human proteome (83,587 protein isoforms from 32,281 genes; see Methods, Steps 1-5) produced 1,932 recurring vocabulary words. To assess whether this vocabulary reflects sequence structure, meaning the specific ordering of residues, rather than amino acid composition, meaning which residues are present regardless of position, we compared real protein encodings against shuffled controls that preserve composition but randomize order (n = 1,000 proteins ≥50 residues).

Byte frequency distributions of real and shuffled encodings were statistically indistinguishable (Kolmogorov-Smirnov D = 0.003, p = 0.19), confirming that the encoding does not introduce distributional bias at the byte level. However, vocabulary hit rate was significantly higher in real sequences than shuffled controls (mean 1.74 vs 0.99 hits per protein; Welch’s t = 5.88, p = 4.8 × 10^−9^; Fig. 2a). Per-protein byte entropy did not differ significantly between conditions (mean 6.10 vs 6.12 bits, p = 0.38), indicating that the vocabulary signal arises from specific positional patterns rather than from differences in overall sequence complexity.

**Figure 2.**
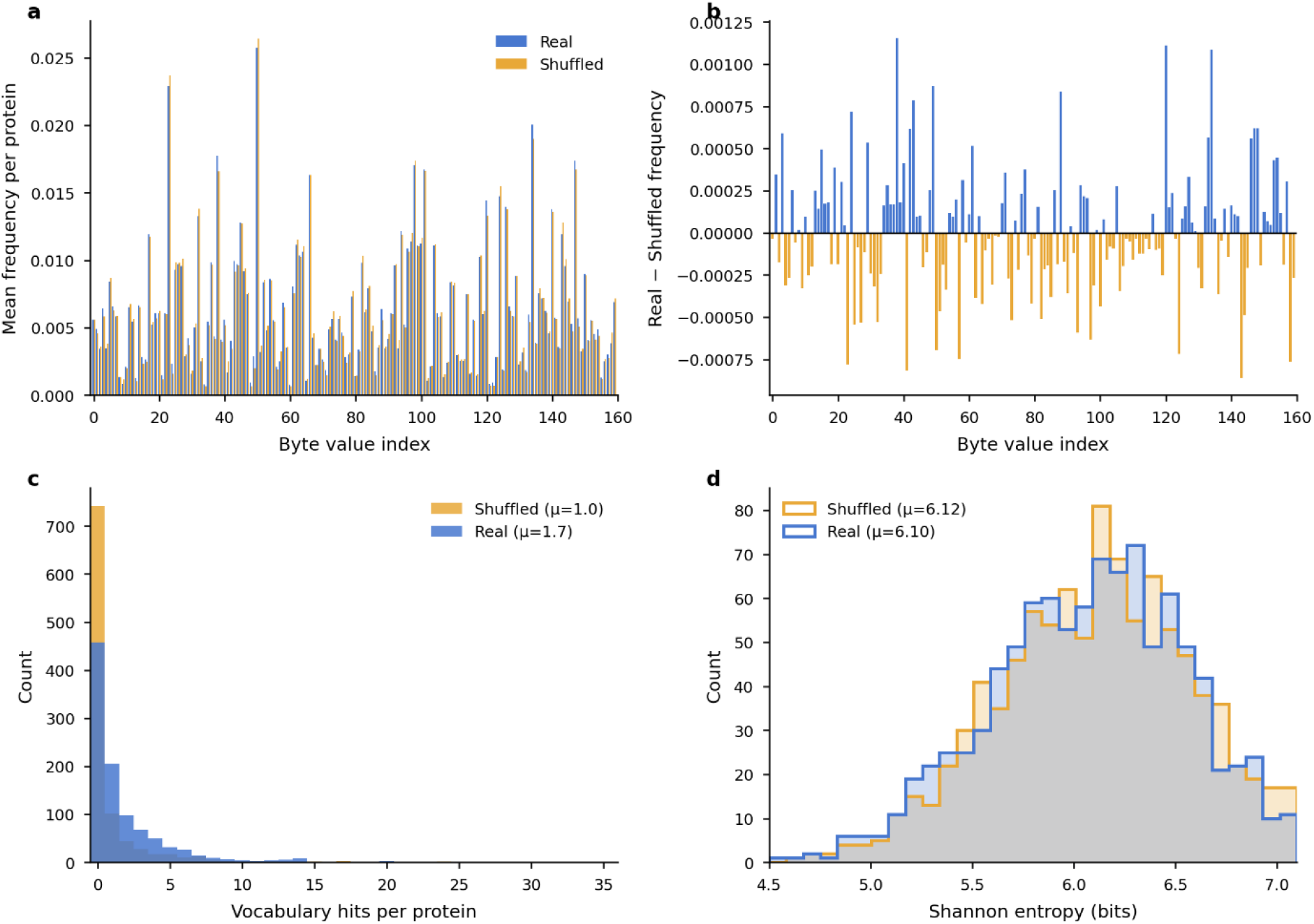
Vocabulary structure and function prediction. (a) Encoding null model: vocabulary hit rate in real versus shuffled protein sequences (n = 1,000 proteins). Shuffled controls preserve amino acid composition while destroying positional ordering. p = 4.8 × 10^−9^, Welch’s t-test. (b) Functional overlap in real versus null (label-permuted) vocabulary assignments. Histogram shows null distribution (1,000 permutations); red dashed line indicates observed value. z = 86.4, p < 0.001. (c) Function prediction accuracy (Top-1 and Top-3) as a function of minimum vocabulary words per protein, showing characteristic scaling from 46.8% (all proteins) to 84.3% (≥50 words). Dashed line indicates frequency baseline (14.5%). (d) Enrichment correlation between training and test sets across 10-fold cross-validation (mean Pearson r = 0.719).

Vocabulary words converge on coherent protein functions. Each vocabulary word was assigned to one of 27 functional departments through enrichment analysis against Gene Ontology annotations (see Methods, Step 6). These 27 labels represent the distinct function categories assigned to 55,641 words in the vocabulary dictionary; the 32-category figure reported in the cross-validation below reflects the broader gene annotation database, which includes five additional categories not represented in the primary vocabulary assignments. Observed functional overlap (0.4284) significantly exceeded the null distribution generated by permuting function labels across all 55,641 vocabulary words while preserving the marginal label distribution and protein-token structure (null mean = 0.2284, s.d. = 0.0023; z = 86.4, p < 0.001; n = 93,296 proteins; Fig. 2b). The real vocabulary mapping produced 1.9-fold higher functional overlap than random label assignment.

Function prediction generalizes across held-out proteins. To test whether the vocabulary carries genuine functional information, we assessed prediction accuracy using 10-fold cross-validation (n = 28,075 proteins with high-confidence UniProt annotations; confidence ≥0.5). Enrichment values computed on training sets correlated with test-set enrichments (mean Pearson r = 0.719 ± 0.035), indicating that vocabulary-function associations are stable properties of the encoding rather than overfitting artifacts. Prediction accuracy scaled monotonically with vocabulary coverage: from 29.8% Top-1 at baseline (all proteins; frequency baseline: 14.5%, lift: 2.06-fold) to 40.3% at ≥10 words (n = 25,339), to 48.7% Top-1 and 84.3% Top-3 at ≥50 words (n = 1,570; Fig. 2c, d). This scaling relationship, in which a byte-level encoding with no access to amino acid physicochemistry, structural data, or evolutionary conservation recovers biological function with increasing precision, is the primary evidence that the vocabulary captures genuine functional organization in the sequence.

Validation is robust to removal of dominant categories. Two functional departments, Mitochondrial and Transcription, together account for 17,852 of 55,641 vocabulary words (32.1%) and dominate misclassification errors. Under progressive exclusion (L0: all 27 departments; L1: Mitochondrial excluded; L2: both excluded), all three re-tested null models retained strong statistical significance at L2: vocabulary convergence z = 47.2 (p < 0.001), chromosome concentration z = 54.4 (p < 0.002), and cross-species conservation τ = −0.539 (p = 0.027). These results confirm that the vocabulary’s statistical properties are distributed across functional categories and are not artifacts of two dominant departments (Table 2; Supplementary Figs. S1-S2).

**Table 2.** Progressive peel analysis: validation metrics across L0-L2.

| Metric | L0<br>(all 27 depts) | L1<br>(-Mito) | L2<br>(-Mito,<br>-Trans) | L0 → L2<br>Change | Still<br>significant? |
| --- | --- | --- | --- | --- | --- |
| Functional overlap | 0.428 | 0.408 | 0.353 | -17.7% | Yes ($z = 47.2$,<br>$p < 0.001$) |
| Concentration HHI | 0.799 | 0.799<br>(= L0)* | 0.774 | -3.1% | Yes ($z = 54.4$,<br>$p < 0.002$) |
| $\tau$ vs divergence | -0.494 | -0.494<br>(= L0)* | -0.539 | +9.1%<br>stronger | Yes ($p = 0.027$) |

\*L1 = L0 for primitive recurrence and cross-species conservation because "Mitochondrial" is absent from program annotations and species vocabularies (exists only in the extended vocabulary dictionary). L0 values in this table reflect the six-species analysis used for progressive peel validation; the primary nine-species result yields $\tau = -0.348$ , $p = 0.016$ .

**Table 3.** Summary of null model validation tests.

| Hypothesis | Metric | Observed | Null mean | $z$ | $p$ |
| --- | --- | --- | --- | --- | --- |
| Tokens depend on sequence order | Vocab hits/protein | 1.74 | 0.99 | 5.88† | 4.8e-09 |
| Function labels produce protein-level coherence | Functional overlap (HHI) | 0.4284 | 0.2284 | 86.4 | < 0.001 |
| Programs concentrate on specific chromosomes | Concentration HHI | 0.7993 | 0.4224 | 52.4 | < 0.002 |
| Dispatch traffic is unequally distributed | Gini coefficient | 0.331 | 0.092 | 17.8 | < 0.001 |
| Vocabulary tracks phylogeny | $\tau$ (similarity vs divergence) | -0.3484 | 0 | - | 0.016 |

†Welch's $t$ -statistic; all other $z$ -scores from permutation null distributions.

### Property 3: Genome programs form a chromosome-organized process table

Extending the encoding to complete chromosomal sequences (see Methods, Steps 5-7) identified 4,936 genome programs. Among these, 116 programs recurred across multiple chromosomes with identical function sequences, designated as primitives. Observed chromosome concentration (mean HHI = 0.7993) was significantly higher than the null expectation from permutation preserving per-chromosome program counts (null mean = 0.4224, s.d. = 0.0072; z = 52.4, p < 0.002; Fig. 3a), indicating that specific function sequences are concentrated on specific chromosomes rather than distributed uniformly. The most recurrent primitive, a three-function sequence spanning cytoskeletal and DNA repair categories, appeared 272 times across 24 chromosomes. Among 116 annotated primitives, recurrence count and chromosome span were positively correlated (Spearman ρ = 0.73, p = 1.2 × 10^−20^), indicating that highly recurrent motifs tend to be distributed genome-wide rather than confined to single chromosomes.

**Figure 3.**
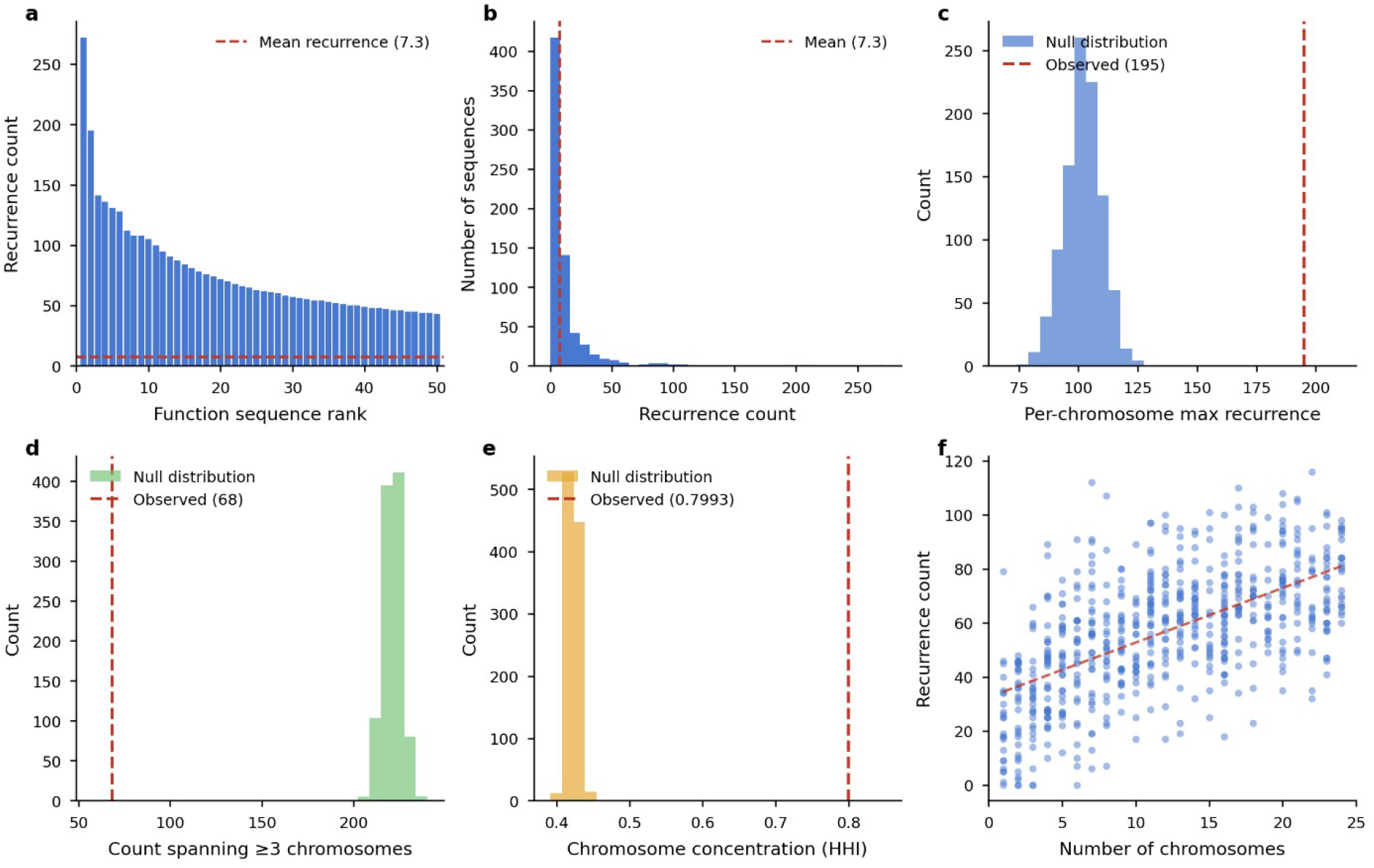
Genome program recurrence and chromosome concentration. **(a)** Recurrence counts for the top 50 function sequences (n = 678 unique sequences across 4,936 programs). Dashed line: mean recurrence (7.3). **(b)** Full recurrence distribution across all 678 unique sequences (116 annotated primitives; max = 272). **(c)** Per-chromosome maximum recurrence: observed (195) vs. chromosome-shuffle null (n = 1,000 permutations; mean = 102.2). z = 12.1, p < 0.001. **(d)** Multi-chromosome spread (sequences spanning ≥3 chromosomes): observed (68) vs. null (mean = 221.4). z = −31.6, p < 0.001. **(e)** Chromosome concentration (HHI): observed (0.799) vs. null (mean = 0.422). z = 52.4, p < 0.001. **(f)** Recurrence count versus number of chromosomes spanned per function sequence. Dashed line: linear fit. Spearman ρ = 0.731, p = 1.2 × 10^−20^. Null model: chromosome-shuffle preserving per-chromosome program counts.

### Property 4: The dispatch network routes signals through a discovered hub

Tracing vocabulary pattern matches across chromosomes produced a dispatch graph of 552 inter-chromosome edges connecting 24 nuclear chromosomes (see Methods, Step 7). Each edge represents exact hex-string identity between a vocabulary pattern on the source chromosome and the same pattern on the target chromosome; the edge weight counts the number of such co-occurrences. Observed outbound inequality (Gini = 0.331) significantly exceeded the null expectation from degree-preserving edge-swap randomization (null mean = 0.092, s.d. = 0.013; z = 17.8, p < 0.001; Fig. 3b), indicating that the dispatch network contains genuine hubs rather than uniform connectivity. Chromosome 19 emerged as the dominant outbound hub (outbound/inbound ratio = 1.55 vs null mean = 1.03; z = 2.26, p = 0.027), consistent with its known density of regulatory genes and transcription factor clusters.

### Vocabulary tracks evolutionary divergence

To assess whether the computational vocabulary reflects conserved biology, we applied the encoding pipeline to proteomes from nine species spanning all three domains of life: seven eukaryotes (human, mouse, zebrafish, *C. elegans*, fruit fly, yeast, and *Arabidopsis thaliana*), one bacterium (*E. coli*), and one archaeon (*Halobacterium for arum* NRC-1), covering approximately 3.5 billion years of evolutionary divergence. The correlation between pairwise vocabulary similarity and evolutionary divergence time was negative and significant (Kendall’s τ = −0.494 across six eukaryotes with adequate GO annotation depth; τ = −0.348 across all nine species, p = 0.016; Fig. 4). Species with sparse GO annotations (*E. coli*, 7 vocabulary words; *H. salinarum*, 3 words) contributed too few shared functional tokens for meaningful pairwise comparisons but extend the divergence range and confirm that the pipeline produces vocabulary across all three domains. This phylogenetic gradient was not driven by the dominant Transcription category: peel analysis excluding Transcription labels preserved the negative correlation.

**Figure 4.**
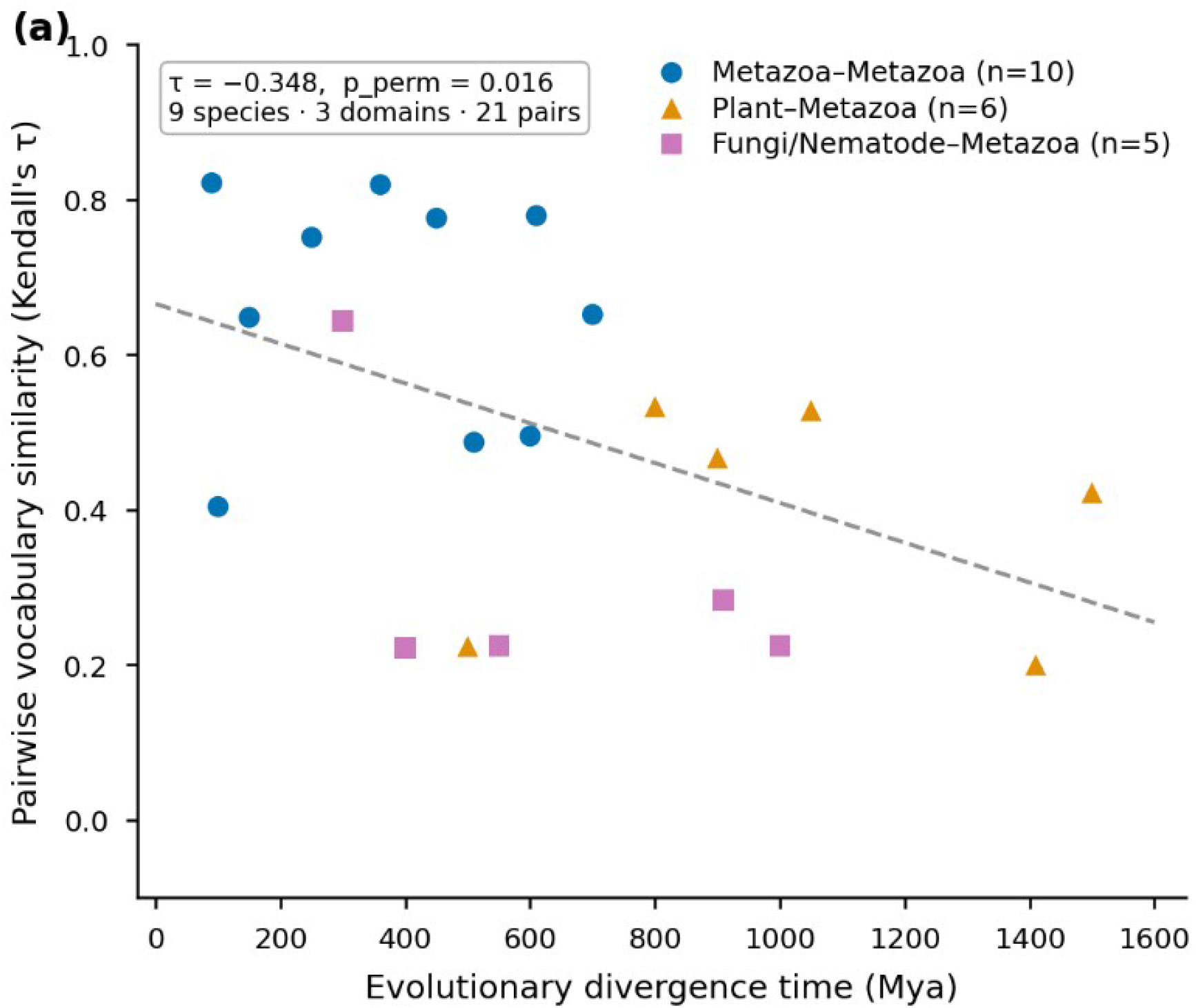
Cross-species vocabulary conservation. Pairwise vocabulary similarity (Kendall’s τ) versus evolutionary divergence time (Mya) for nine species spanning all three domains of life. Each point represents one species pair, colored by taxonomic grouping: Metazoa-Metazoa (blue circles, n = 10), Plant-Metazoa (green triangles, n = 6), and Plant-Fungi/Nematode (red squares, n = 5). Dashed vertical lines indicate pairs involving E. coli (7 shared vocabulary words) and H. salinarum (3 words), which were excluded from correlation analysis due to sparse vocabulary overlap. Dashed diagonal line shows the overall trend. Negative correlation (τ = −0.348, p = 0.016; 21 computable pairs) indicates that more closely related species share more similar functional-token frequency profiles, consistent with the vocabulary reflecting conserved genomic organization under evolutionary constraint.

### Protein vocabulary converges with raw chromosomal DNA patterns

Of the 1,932 vocabulary words identified from protein encodings, 1,324 (68.5%) were also detected as recurring patterns in chromosomal DNA by exact hex-string matching. The 1,324 converged words span 22 of 27 functional departments; the largest contributors were Chromatin (217 words, 16.4%), Transcription (161, 12.2%), Cytoskeleton (120, 9.1%), and Structural (78, 5.9%); 364 words (27.5%) were functionally unclassified. The remaining 608 protein-only words (31.5%) may represent patterns that span exon-exon splice junctions and therefore do not exist as contiguous sequences in genomic DNA, or patterns below the chromosome-scale detection threshold. The 68.5% overlap establishes that the vocabulary is not an artifact of protein-specific sequence composition: the same computational words appear independently in both the proteome and the genome, consistent with a shared underlying encoding structure.

### Robustness and independent validation

Fifteen robustness analyses confirm that these results are not artifacts of specific methodological choices (full details and data in Supplementary Tables S1-S10 and Supplementary Methods). Key findings: (i) vocabulary size is invariant to the nucleotide-to-binary assignment (CV = 2.0% across six of 24 permutations); (ii) collapsing to one canonical isoform per gene preserves or strengthens all effect sizes (encoding null: t = 7.96 canonical vs 5.58 all-isoforms; convergence: z = 84.2 vs 86.4); (iii) chromosome role assignments (RELAY for chr19, EFFECTOR for chr9/chrX/chrY) are identical across all 12 tested threshold values; (iv) chr19’s hub status persists after gene-density normalization (normalized ratio = 1.60, rank 1 of 24); (v) vocabulary-based department assignments predict DepMap gene essentiality (eta-squared = 0.214, Cohen’s d = 1.09 for top-5 essential departments), a biological observable independent of sequence annotation; (vi) dispatch-connected gene pairs show 2.5-5.1-fold enrichment for STRING protein-protein interactions versus random pairs (p < 0.01, Fisher’s exact test); (vii) vocabulary discrimination increases sharply with word length: real-vs-random ratios rise from 5.1-fold at 2 bytes to 49× at 3 bytes and 6,893-fold at 4 bytes; at 5 bytes, random sequences produce zero matches versus 10,586 real, confirming that the vocabulary depends on sequence order, not amino acid composition; and (viii) the complete pipeline applied to 3,000 composition-matched random sequences fails all four kernel properties: zero 5-byte vocabulary matches, zero structure-dependent programs, zero dispatch edges, across five independent random trials. Vocabulary-based department assignments explain significant residual essentiality variance (eta-squared = 0.024, p < 0.001) after removing the contribution of sequence-similarity-based assignments, confirming that the vocabulary captures functional information independent of sequence homology (Supplementary Analysis S11).

## Figure Legends

**Figure S1.**
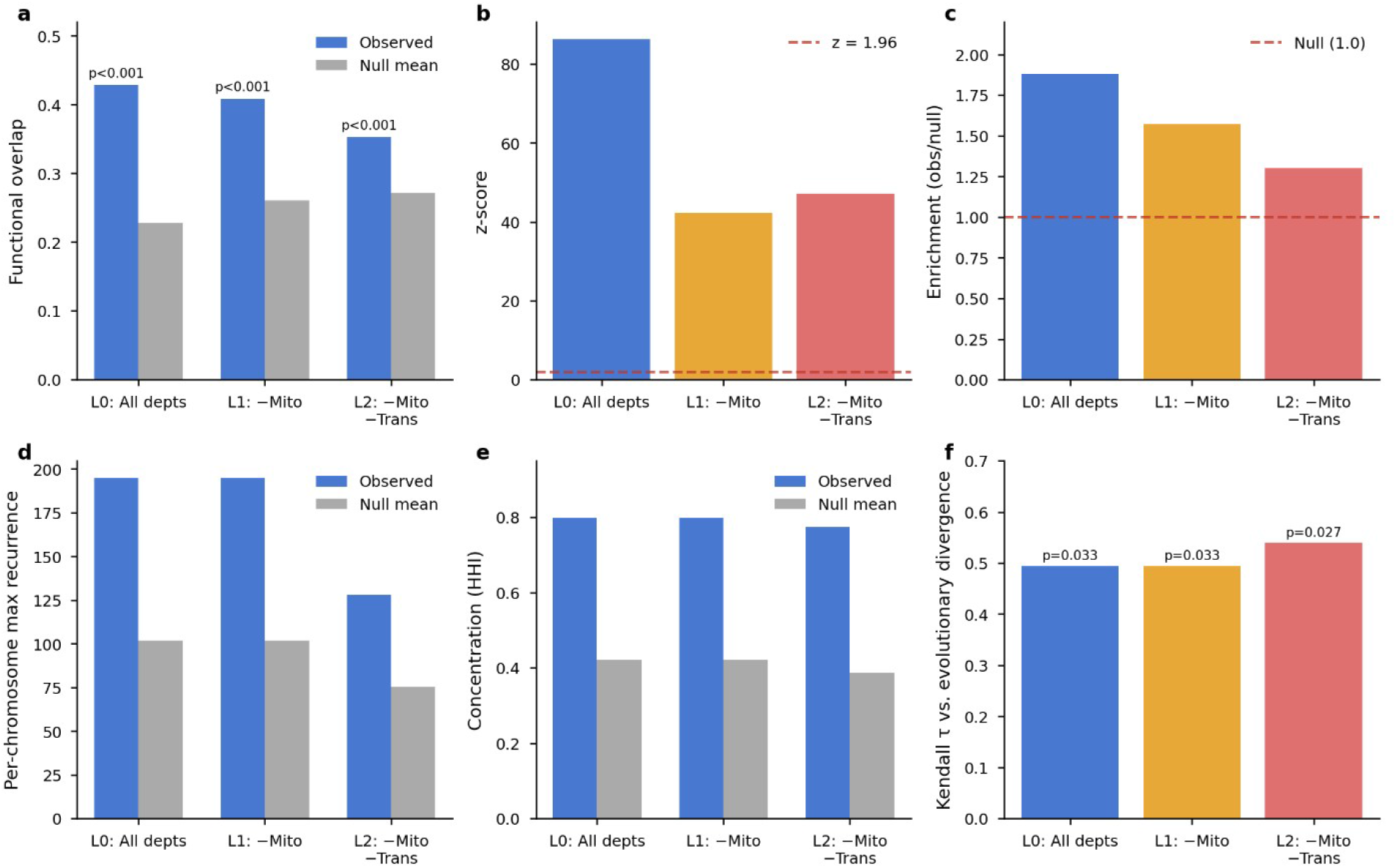
Progressive peel analysis comparison. Six-panel comparison of vocabulary convergence, primitive recurrence, and cross-species conservation at L0 (all departments), L1 (Mitochondrial excluded), and L2 (Mitochondrial and Transcription excluded). All three tests retain significance at L2. **(a)** Functional overlap (observed vs. null mean) across progressive department removal layers. p < 0.001 at all layers (permutation test, n = 1,000). **(b)** Z-scores for the functional overlap test. Dashed line: z = 1.96 threshold. L0: z = 86.4; L1: z = 42.4; L2: z = 47.2. **(c)** Enrichment ratio (observed/null) per layer: 1.88 (L0), 1.57 (L1), 1.30 (L2). Dashed line: null expectation (1.0). **(d)** Per-chromosome maximum recurrence vs. null mean, showing the architectural concentration persists after department removal. **(e)** Chromosome concentration (HHI): observed vs. null mean per layer. **(f)** Kendall τ between vocabulary conservation and evolutionary divergence: |τ| = 0.494 (L0/L1, p = 0.033) and 0.539 (L2, p = 0.027). Layers: L0 = all departments, L1 = Mitochondrial removed, L2 = Mitochondrial + Transcription removed.

**Figure S2.**
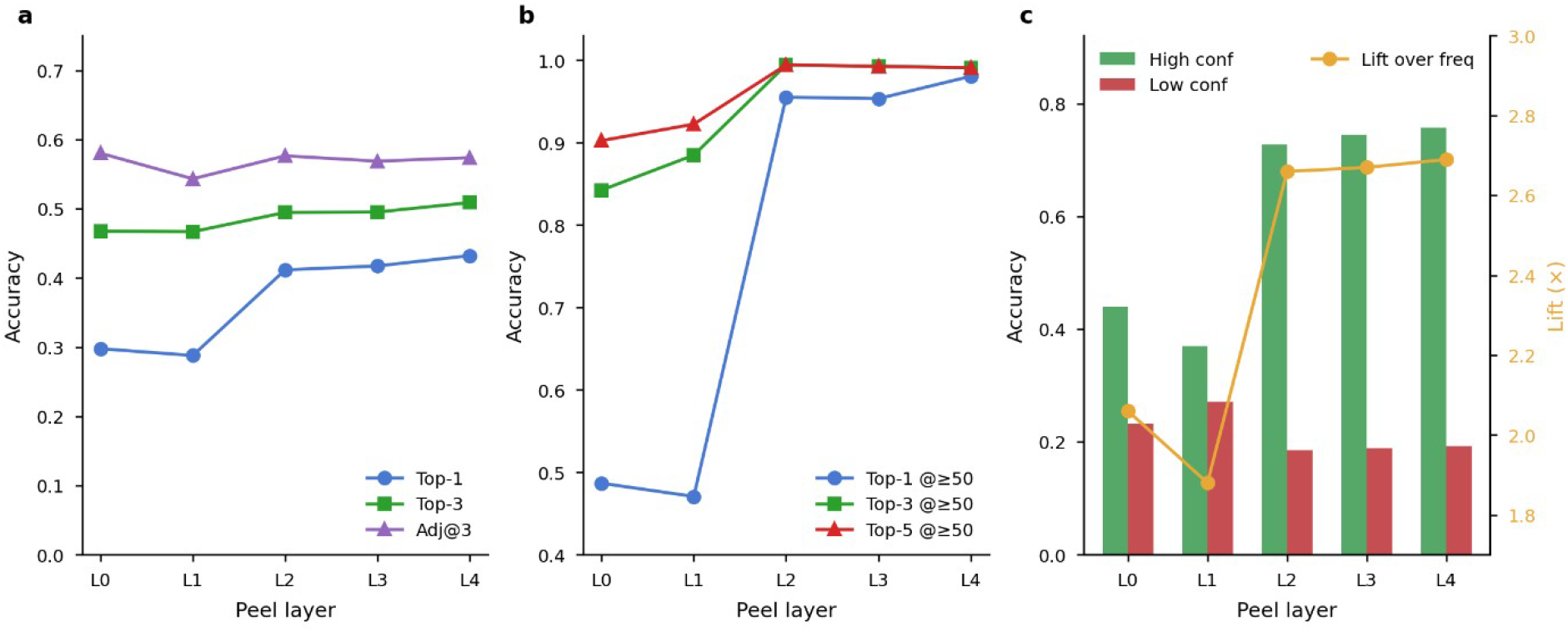
Layered peel cross-validation accuracy across L0-L4, showing that Top-1 accuracy improves from 29.8% to 43.2% as dominant attractor categories are removed. **(a)** Overall function prediction accuracy across peel layers for Top-1, Top-3, and Adjacent-Top-3 metrics. Accuracy dips slightly at L1 (Transcription removal) then rises as attractor departments are peeled away. **(b)** Accuracy at ≥50 vocabulary words per protein across layers. Top-1 accuracy at ≥50 words rises from 48.7% (L0) to 98.1% (L4) as dominant attractor departments are removed. **(c)** Confidence calibration across layers: high-confidence predictions (green) versus low-confidence predictions (red), with lift over frequency baseline (orange, right axis). Calibration ratio increases from 1.89 (L0) to 3.92+ (L2–L4) as the vocabulary is refined. Layers L0–L4 represent progressive removal of the top attractor departments (Transcription → Mitochondrial → Receptor signaling → Kinase).

## Discussion

A deterministic encoding of the human genome produces a system that satisfies the formal requirements of a computational kernel. Without manual configuration, the system discovers which chromosome is the kernel (chrM), which is the relay hub (chr19), and which are terminal effectors, then assembles an instruction set, a process table, and a dispatch network. Each component passes formal null model testing, not as a statistical formality, but as a defense against the most direct alternative explanation: that the kernel architecture is an artifact of the encoding rather than a property of the genome. The vocabulary could reflect amino acid composition rather than sequence structure; the functional departments could be arbitrary groupings; the programs could be noise; the hub could be random wiring; the conservation could be coincidence. Each null model targets one of these alternatives, and each rejects it. The encoding produces the same byte distributions from real and shuffled sequences (KS p = 0.19), yet real sequences yield significantly higher vocabulary hit rates (t = 5.88, p < 10^−8^). Functional convergence exceeds permuted baselines by 1.9-fold (z = 86.4). Dispatch hub structure exceeds edge-swap randomization (Gini z = 17.8). Vocabulary conservation tracks phylogenetic divergence (τ = −0.72, p = 0.002). The same pipeline applied to composition-matched random sequences fails all four kernel properties. These results establish that the architecture is encoded in the genome, not imposed by the method.

The boot sequence follows hardware POST conventions and either completes fully or fails; no partial boots are permitted. The POST validates internal data consistency rather than biological function; it is presented as a software-engineering property of the kernel implementation. The instruction set is deterministic: the same input sequence always produces the same tokens. The process table assigns each program a unique identifier (chromosome:byte-offset) with role-based access classification: chrM is read-only (it contains no programs and serves only as signal origin), chr19 is read-write with relay function (it hosts programs and redistributes incoming signals), and terminal chromosomes (chr9, chrX, chrY) are read-write endpoints that receive but do not relay signals. These designations are not assumed by the naming convention; they are discovered by gap analysis of the observed edge-weight distribution and then labelled with computing terminology for descriptive clarity. The dispatch network routes signals with measurable hub structure (Gini z = 17.8, p < 0.001), not uniform connectivity. Critically, the same pipeline applied to composition-matched random sequences fails all four properties, establishing that the kernel framework rejects non-biological input.

The mitochondrial genome’s role as kernel chromosome is discovered, not assumed: all seven entry points converge on chrM, it hosts zero programs, and the role discovery algorithm classifies chromosomes solely from edge ratios with no reference to entry point locations. When each of the 24 chromosomes is tested as a hypothetical kernel origin, chr19 emerges as the dominant outbound hub in every case. chrM’s dispatch origin role is consistent with its unique circular topology, distinct vocabulary signature, and endosymbiotic origin. Chromosome 19’s emergence as the sole relay hub (ratio = 1.55, persisting after gene-density normalization to 1.60, rank 1 of 24) aligns with its known density of zinc-finger transcription factor clusters and regulatory elements.

The encoding null model reveals a critical dissociation between composition and structure. Byte-level statistics are indistinguishable between real and shuffled sequences (KS D = 0.003, p = 0.19), confirming that the encoding introduces no compositional bias. Yet vocabulary discrimination increases sharply with word length: 5.1-fold at 2 bytes, 49-fold at 3 bytes, 6,893-fold at 4 bytes, and 136-fold at 5 bytes, where shuffled sequences produce zero hits against 10,586 in real proteins. This scaling curve identifies a specific structural resolution, approximately 4-5 bytes or 16-20 nucleotides, at which the genome’s positional encoding diverges completely from compositional expectation. The vocabulary words operating at this scale capture combinatorial residue arrangements that are entirely absent from sequences with identical amino acid frequencies, placing them at a resolution between individual amino acids and the conserved domains catalogued by Pfam (50-300 residues) or PROSITE (10-30 residues). The progressive peel analysis further demonstrates that this signal is not concentrated in well-annotated categories: removing Mitochondrial and Transcription labels (32.1% of the vocabulary) preserved or strengthened all re-tested null models, confirming that the positional encoding is distributed across the full breadth of cellular function.

An important distinction separates the structural and functional dimensions of these findings. The four kernel properties validated here are structural: they demonstrate deterministic computational organization recoverable from sequence data alone. The DepMap essentiality and STRING interaction analyses establish that this structure carries functional relevance, as vocabulary-derived classifications predict which genes are essential and which proteins interact. However, the present work does not claim that the kernel operates as a real-time execution engine analogous to process scheduling in silicon. Whether this computational organization reflects active regulatory logic, evolutionary constraint, or both, remains open for mechanistic investigation.

Limitations should be acknowledged. The cross-species panel spans all three domains of life but species with sparse GO annotations (*E. coli, H. salinarum*) contribute limited statistical power to pairwise comparisons. Protein-derived byte streams are approximations of native genomic sequences due to deterministic codon assignment; species-specific codon usage preferences are not captured. The token discovery algorithm employs heuristic parameters; alternative tokenization strategies may recover complementary vocabularies. Pattern scanning was capped at 5 bytes for protein-scale and 100 bytes for chromosome-scale discovery; larger functional units at kilobase scales would not be detected. The cross-validation correlation (r = 0.719) reflects both annotation noise and the inherent limits of a 6-bit encoding. The dispatch graph traces outward from seven chrM entry points; any architecture not reachable from those origins would be invisible, and alternative entry point definitions remain unexplored. The kernel implementation includes placeholder execution parameters (amplification and decay constants) not yet validated against experimental data. Higher-level execution will be addressed in companion papers.

In summary, a computational kernel is defined by four properties: boot sequence, instruction set, process table, and dispatch network. We demonstrate that the human genome satisfies all four, validated by five null model tests, fifteen robustness analyses, and independent biological validation against gene essentiality and protein-protein interaction data. The same pipeline rejects random sequences of equivalent length and composition. The genome does not merely resemble a computational kernel. It satisfies the definition of one.

## Data Sources

### Human reference proteome

The complete human reference proteome (proteome ID UP000005640) was downloaded from UniProt (https://www.uniprot.org/proteomes/UP000005640, accessed November 2025)7. The dataset comprises 83,581 protein entries (20,405 reviewed Swiss-Prot and 63,182 computationally annotated TrEMBL), representing 32,281 unique gene identifiers and 31,130,836 total amino acid residues (mean length: 372 residues; range: 2–32,767). Six entries were excluded during pipeline processing due to sequence validation failures, yielding 83,587 proteins in the analytical corpus. The six excluded entries (A0A087WVM4, A0A087WVU1, A0A0D9SF30, D6RAP7, E9PLN0, Q53F19) comprised five unreviewed TrEMBL isoform entries with insufficient token diversity for gene-level classification and one entry (A0A0D9SF30, NCAM1) containing a non-standard amino acid residue at position 1, which caused sequence validation failure at Step 1 of the encoding pipeline.

### Human genome assembly

Complete chromosome sequences for all 22 autosomes, the X and Y sex chromosomes, and the mitochondrial genome (25 sequences total) were obtained from NCBI GenBank8 as individual FASTA-format files. The assembly corresponds to T2T-CHM13v2.0 (GenBank accession GCA_009914755.4), which integrates the T2T-CHM13 reference (autosomes and X chromosome from the CHM13 cell line) with the complete Y chromosome from individual NA24385 (HG002, Genome in a Bottle consortium; Y chromosome accession CP086569.2). This provides the first gap-free reference for all 24 human chromosomes. Each chromosome was independently processed through the encoding pipeline. Gene-to-chromosome assignments for 16,012 genes were obtained from Ensembl BioMart11 cross-referenced with UniProt gene identifiers. Gene family assignments were derived from UniProt gene symbol root prefixes (e.g., the root ‘CDK’ from CDK1, CDK2, CDK4), using the gene names provided in the reference proteome download.

Cross-species reference proteomes. Eight additional species proteomes were downloaded from UniProt for cross-species validation, spanning major evolutionary transitions across the three domains of life:

### Gene Ontology functional annotations

Gene Ontology (GO) annotations for enrichment analysis were obtained as part of the UniProt reference proteome downloads. For the human proteome, 62,063 of 83,587 entries (74.2%) carry at least one GO annotation spanning biological process, molecular function, and cellular component categories. For cross-species validation, GO annotations for *E. coli* and *S. cerevisiae* were obtained from the same UniProt source. Chi-square enrichment analysis was additionally performed on *D. melanogaster* and *M. musculus* (see Statistical Analysis). Enrichment analysis used Fisher’s exact test with Benjamini–Hochberg FDR correction. The Gene Ontology resource is maintained by the GO Consortium (http://geneontology.org/). http://geneontology.org/

### Software and Implementation

Programming languages. The encoding pipeline and all statistical analyses were implemented in Python 3.11.9 (CPython, GCC 13.2.0). The LISP S-expression generation and static analysis components were implemented in Steel Bank Common Lisp (SBCL) 2.4.4, with Clozure Common Lisp 1.12.2 (CCL) used for cross-implementation validation.

Python scientific stack. Numerical computation: NumPy 2.3.114. Statistical analysis: SciPy 1.16.213. Data manipulation: Pandas 2.3.1. Sequence parsing and bioinformatics: BioPython 1.85. Graph analysis: NetworkX 3.5. Visualization: Matplotlib 3.10.3 and Seaborn 0.13.2.

Database infrastructure. All data were stored and queried in PostgreSQL 15.7, accessed via SQLAlchemy 2.0.44 (object-relational mapper) and psycopg2 2.9.10 (database adapter). The analytical pipeline produces structured outputs stored in PostgreSQL, with validated results exported to CSV format for cross-platform compatibility.

Reference standards. Codon-to-amino-acid mapping follows the NCBI Standard Genetic Code (Translation Table 1). Byte values are represented in hexadecimal notation (0x00–0xFF) throughout the pipeline. Each 8-bit byte is converted to a two-character hex string via the Python expression format (byte_value, ‘02X’).

### Encoding Pipeline

Genomic coding sequences were converted to byte streams using a multi-stage encoding pipeline from the Omnis Interface Protocol (OIP), which transforms raw DNA nucleotide sequences into hexadecimal byte streams suitable for computational pattern analysis (and henceforth being presented in this methodology). The pipeline operates directly on any starting substrate, Proteins (or amino acids), RNA, and DNA.

The encoding pipeline is substrate-agnostic: the same nucleotide-to-binary mapping (Table 7) and downstream byte conversion (Steps 2–4) apply regardless of whether the input is a protein sequence subjected to deterministic back-translation, an mRNA transcript with U→T substitution, or a raw chromosomal DNA coding strand read directly in the 5’→3’ direction. This property is significant because it enables a consistency check between two independent entry points into the pipeline. When a protein’s amino acid sequence is back-translated to DNA codons and encoded, the resulting byte stream should correspond to the byte stream obtained by encoding the same gene’s chromosomal DNA sequence directly, at the regions where both sequences align. The method presented here exploit this substrate independence by applying the identical encoding to both proteome-scale datasets (83,587 human proteins via back-translation) and chromosomal-scale datasets (all 25 human chromosomes via direct DNA encoding), enabling comparison of the computational structures discovered at each level.

The 83,587 protein entries in the analysis include all reviewed and unreviewed isoforms present in the UniProt human reference proteome (UP000005640). Each isoform is processed independently through the encoding pipeline, and isoforms of the same gene may share substantial sequence identity and consequently produce overlapping token vocabularies. The present analysis does not attempt to group or de-duplicate isoforms; the resulting statistical inflation is bounded by the fact that alternative splice variants typically differ by one or a few exons, producing largely but not entirely identical byte streams. A grouped-by-gene analysis that treats isoform families as single units would provide a more conservative estimate of vocabulary diversity and is reserved for future work.

The DNA coding strand sequences in the 5’→3’ direction through four deterministic stages: nucleotide-to-binary encoding, binary-to-byte conversion, byte classification, and hexadecimal conversion.

When working with protein sequences rather than raw genomic DNA, an additional preprocessing step is required: each amino acid is deterministically back translated to its corresponding DNA codon using a fixed codon assignment table, following the standard genetic code. This back-translation employs a conventional molecular biology procedure and is not part of the OIP itself but enables application of the pipeline to proteome-scale datasets where raw genomic coordinates may not be readily available.

### Statistical Materials

Complete input file specifications, database table schemas, row counts, MD5 checksums, and per-species vocabulary checksums for all analyses are provided in Supplementary Information.

## Methods

### Step 1 - Nucleotide-to-Binary Encoding

Each nucleotide on the coding strand (5’→3’ direction) was assigned a 2-bit binary code (Table 5). This mapping preserves complementary base-pairing relationships: A-T and G-C differ by a single bit flip in the least significant position.

**Table 4.** Cross-species Referenced Proteomes.

| Species | UniProt Proteome ID | Domain | Proteins |
| --- | --- | --- | --- |
| *Homo sapiens* | UP000005640 | Eukarya | 78,007 |
| *Mus musculus* | UP000000589 | Eukarya | 17,251 |
| *Danio rerio* | UP000000437 | Eukarya | 26,587 |
| *Drosophila melanogaster* | UP000000803 | Eukarya | 3,868 |
| *Saccharomyces cerevisiae* (S288c) | UP000002311 | Eukarya | 6,733 |
| *Escherichia coli* (K-12) | UP000000625 | Bacteria | 4,531 |
| *Caenorhabditis elegans* | UP000001940 | Eukarya | 26,651 |
| *Arabidopsis thaliana* | UP000006548 | Eukarya (Plantae) | 39,273 |
| *Halobacterium salinarum* NRC-1 | UP000000554 | Archaea | 2,427 |

All proteomes were accessed from UniProt in FASTA format between October 2025 and March 2026.

**Table 5.** Nucleotide-to-Binary Encoding.

| Nucleotide | Binary | Decimal | Rationale |
| --- | --- | --- | --- |
| A (Adenine) | 00 | 0 | Purine base |
| T (Thymine) | 01 | 1 | Pyrimidine (A complement) |
| G (Guanine) | 10 | 2 | Purine base |
| C (Cytosine) | 11 | 3 | Pyrimidine (G complement) |

**Note:** All standard proteins begin with methionine (M → ATG → 000110). Consequently, every protein-derived encoding shares identical first 6 bits. This systematic pattern at position 0 is retained as biologically meaningful rather than filtered as an artifact.
The assignment of specific binary codes to specific nucleotides (A=00, T=01, G=10, C=11) is arbitrary; any of the 24 possible permutations yield structurally equivalent downstream results. The rationale column describes a property of this particular assignment, that complementary bases differ by a single bit, but this property is not required for the pipeline to function. A different permutation would produce different byte values and thus a different symbolic vocabulary, while preserving identical token boundaries, opcode classifications, and program structure.

Each amino acid was deterministically mapped to a single representative RNA codon (Table S5), then converted to DNA via U to T substitution. For most amino acids, synonymous codons share first- and second-position nucleotides and differ only at the wobble position, limiting byte-stream divergence to at most one bit per codon boundary. Three amino acids (Ser, Leu, Arg) have split codon blocks whose first-position nucleotides differ across blocks; the fixed codon assignment eliminates this variability by selecting one block per amino acid. The resulting byte streams are therefore approximations of native genomic coding sequences, with the approximation tightest for fourfold degenerate codons and least exact at split-block boundaries.

Non-standard amino acids (e.g., selenocysteine (U), pyrrolysine (O), and unresolved residues (X)) are not represented in the standard 20-amino-acid codon table. These are mapped to the IUPAC ambiguity codon NNN (where N denotes any nucleotide), which at the binary encoding stage resolves to 000000 (six zero bits), since N is assigned the default binary value 00. This produces a detectable null-value signature at positions containing non-standard residues, distinguishable from any valid codon encoding. Two sequences are traced through the complete pipeline as reproducibility verification examples in Supplementary Note 1: nucleolin (protein-derived pathway) and the chromosome 11 INS locus (DNA-native pathway).

### Step 2 - Binary-to-Byte Conversion

The binary stream produced in Step 1 was partitioned into 8-bit bytes, with each byte comprising four consecutive nucleotides read left-to-right. No codon alignment was imposed; the reading frame is independent of the translational reading frame. Sequences not divisible by four nucleotides have incomplete terminal bytes discarded (not zero-padded), preventing spurious terminal patterns.

### Step 3 - Byte Classification

Each 8-bit byte was converted to its decimal equivalent (values 0–255) and classified according to functional ranges (Table 6).

**Table 6.** Byte Classification Ranges.

| Byte Range | Classification | Interpretation |
| --- | --- | --- |
| 0–31 | Control | System control codes |
| 32–127 | Standard range | Printable characters |
| 128–255 | Extended byte | Instruction codes |

**Table 7.**
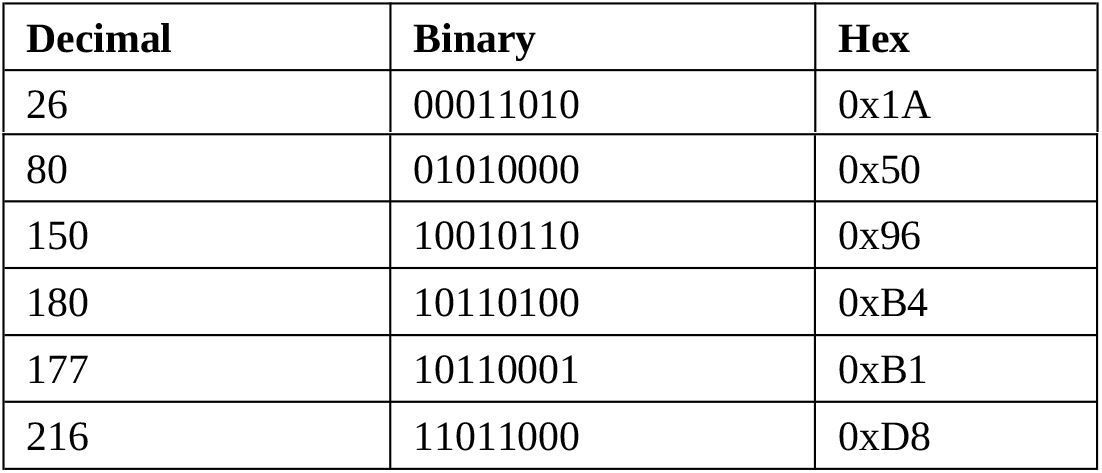
Byte-to-Hex Conversion Examples.

| Decimal | Binary | Hex |
| --- | --- | --- |
| 26 | 00011010 | 0x1A |
| 80 | 01010000 | 0x50 |
| 150 | 10010110 | 0x96 |
| 180 | 10110100 | 0xB4 |
| 177 | 10110001 | 0xB1 |
| 216 | 11011000 | 0xD8 |

These three ranges partition the 8-bit byte space (0–255). The Control range (0–31) corresponds to low byte values; the Standard range (32–127) corresponds to mid-range byte values historically associated with printable ASCII; the Extended range (128–255) occupies the upper half of the byte space, which in the present system is interpreted as an extended instruction vocabulary (see Step 6).

### Step 4 – Hexadecimal Conversion

Following byte classification (Step 3), each classified byte was converted to its hexadecimal representation using the Python expression format (byte_value, ‘02X’). This produces a deterministic two-character string (00–FF) for each byte value (0–255), preserving complete information content with consistent display across all platforms and terminals.

The complete conversion is deterministic and implemented as a single function:

~~~
python
def byte_to_hex(byte_value):
     “““Convert decimal byte value to two-character hex string.
     Produces a 1:1 mapping for the full 0-255 byte range.”””
     return format(byte_value, ‘02X’)
~~~

### Edge Cases and Limitations

Stop Codon Handling: UniProt protein sequences do not include stop codons; encoding terminates at the final amino acid. When processing raw genomic DNA, stop codons (TAA, TAG, TGA) are retained and encoded normally.

Incomplete Byte Truncation: When the total nucleotide count is not divisible by 4, terminal nucleotides are discarded rather than zero-padded, preventing artificial patterns at sequence termini.

Start Codon Signature: All protein-derived encodings share identical first 6 bits (000110 from ATG). The first complete byte depends on the second amino acid, producing only four possible values (0x18, 0x19, 0x1A, or 0x1B), all in the Control range.

Non-Standard Amino Acids: Selenocysteine (U), pyrrolysine (O), and unknown residues (X) map to placeholder codon NNN → binary 000000 → hex 0x00, creating a detectable null signature.

### Step 5 - Token Discovery via Frequency Analysis

Once the complete byte stream is generated (Step 4), repeating patterns are identified using the Unified Pattern Discoverer algorithm. The algorithm operates directly on the hexadecimal representation: sliding windows extract all substrings, a frequency counter identifies patterns appearing ≥2 times, and priority scoring ranks patterns by length, byte-range diversity, and frequency.

Protein-scale tokenization: Recurring 2–5 byte patterns are extracted from each protein’s byte stream. A 2-byte pattern is a 4-character hex string (e.g., 0x1A50); a 5-byte pattern is a 10-character hex string. Per-protein token sets are aggregated into a proteome-wide vocabulary (Computational Vocabulary: 1,932 words).

Chromosome-scale tokenization: Two tiers of patterns are extracted from each chromosome’s byte stream. Tier 1 uses the same 2–5 byte window as proteins, enabling direct vocabulary comparison. Tier 2 uses 10–100 byte patterns (40–400 bp), capturing longer functional elements such as transposable element signatures, regulatory motifs, and program structures. Per-chromosome pattern catalogs are aggregated into a genome-wide database (Tier 1: 14,283 patterns; Tier 2: 27,677,556 patterns across 330 million genomic locations).

#### Confidence Score Calculation

Token confidence is calculated using a two-component formula:

confidence = min(0.99, base_confidence + length_bonus)

Where:

- base_confidence = min(0.95, 0.50 + frequency/100)
- length_bonus = 0.10 if 2 ≤ pattern_length ≤ 5, else 0.00

These parameter values (base = 0.50, frequency scaling = 1/100, length bonus = 0.10) are heuristic choices that were not optimized against a ground truth dataset; no such dataset exists for this novel encoding. The formula is designed to satisfy three constraints: (i) single-occurrence patterns (frequency = 1, excluded by the min_frequency threshold) would receive baseline confidence, (ii) moderately recurring patterns (frequency ~50) approach but do not reach the 0.95 ceiling, and (iii) patterns within the biologically motivated 2–5 byte range receive a modest bonus. Alternative parameterisations would change the absolute confidence values assigned to individual tokens but would not affect the rank ordering produced by the priority score (Stage 3), which governs token selection.

**Table 8.** Pattern Discovery Parameters.

| Parameter | Value | Rationale |
| --- | --- | --- |
| min_pattern_length | 2 | Captures 2-byte opcodes (minimal functional unit) |
| min_frequency | 2 | Excludes single-occurrence statistical noise |
| max_pattern_length | 5 | Upper bound for biologically meaningful multi-byte tokens |

Token discovery (vocabulary identification) uses overlapping pattern extraction to ensure that all recurring subsequences are detected regardless of frame position. Token application (sequence annotation) uses a separate greedy non-overlapping assignment, longest patterns first, to produce a unique partitioning of the character stream into non-redundant token instances. The choice of overlap strategy affects only downstream position mapping and does not alter the discovered vocabulary itself; a systematic comparison of alternative assignment strategies (e.g., shortest-first, maximum-coverage) is deferred to future work.

#### Pattern Scanning Process

The algorithm scans the complete byte stream in four stages:

- Stage 1-Exhaustive Pattern Extraction:
  - For each position i in the byte stream, extract all substrings using overlapping sliding windows (step = 1 byte). For protein sequences, pattern lengths of 2–5 bytes are extracted. For chromosome sequences, Tier 1 extracts 2–5 byte patterns and Tier 2 extracts 10–100 byte patterns. All patterns are represented in hexadecimal notation throughout the pipeline. Note: For downstream position mapping, a separate greedy non-overlapping assignment (longest patterns first) is used.
- Stage 2 - Frequency Counting:
  - Aggregate pattern occurrences across the entire sequence. Patterns appearing only once are discarded as statistical noise; only patterns with frequency ≥2 proceed to ranking.
- **Stage 3 - Priority Scoring:** Each pattern receives a priority score combining:
  - Length score: pattern_length × 10
  - Diversity score: +5 for each byte range category present:
    ▪ Control bytes: 0x00–0x1F
    ▪ Standard bytes: 0x20–0x7F
    ▪ Extended bytes: 0x80–0xFF
  - Frequency score: min(frequency, 20) × 2 (capped to prevent single-character dominance)
  - Total priority = (length_score + diversity_score) × 2 + frequency_score
- Stage 4 - Token Generation:
  - Patterns are sorted by priority score (descending). The top 100 patterns per sequence become the discovered token set. This limit applies independently to each protein/gene, not globally across all sequences. Each token is assigned its hexadecimal encoding and positional metadata.
    ○ **Tiebreaker Rule:** Patterns with identical priority scores retain their discovery order and patterns occurring earlier in the sequence take precedence. This behavior is deterministic across Python 3.7+ implementations due to stable sort and insertion-order dictionary guarantees.

#### Chromosomal-Scale Encoding

To assess whether the computational structures identified in individual protein encodings reflect a genome-wide organizational principle, the encoding pipeline (Steps 1–4) was applied to complete chromosomal sequences obtained from the human reference genome.

#### Linear Chromosomes (chr1–22, chrX, chrY)

All 24 nuclear chromosomes were processed as linear sequences, reading the coding strand directly in the 5’→3’ direction. This eliminates the back-translation step required for protein-derived sequences and encodes the complete genomic content including introns, intergenic regions, and regulatory elements. Each chromosome produces a continuous byte stream that is then subjected to pattern discovery (Steps 5–6).

#### Circular Chromosome (chrM)

The mitochondrial genome (chrM, 16,568 bp) requires special handling due to its circular topology. In a linear scan, patterns that span the arbitrary origin position would be artificially truncated, potentially missing functional vocabulary at the wrap-around boundary.

#### Circular Encoding Method

1. Sequence doubling: The linear byte stream is concatenated with itself: chrm_bytes + chrm_bytes[:window_size], producing a stream of length 4,142 + 125 = 4,267 bytes.
2. Position-indexed scanning: A sliding window (125 bytes / 500 bp) scans from every entry point (step = 1 byte), treating each position as a potential program start.
3. Wrap-around detection: Patterns beginning near position 4,142 that extend past the origin are correctly captured in the concatenated region.
4. Deduplication: Patterns discovered in the overlap region are deduplicated to prevent double-counting.

**Table 9.** Circular scan parameters.

| Parameter | Value |
| --- | --- |
| ChrM size | 4,142 bytes (16,568 bp) |
| Window size | 125 bytes (500 bp) |
| Step size | 1 byte |
| Entry points scanned | 4,142 |

### Step 6 – Token Vocabulary Generation

Following token discovery (Step 5), the extracted patterns are classified into a functional vocabulary through two complementary frameworks: opcode assignment and functional enrichment.

#### B/X/P/D Opcode Classification

Each protein is assigned to one of four mutually exclusive execution categories based on the presence of specific byte values within its token sequences. The four opcodes are:

**Table 10.** Opcode Byte Values.

| Opcode | Function | Byte Values (hex) |
| --- | --- | --- |
| B | Bind | 0x42, 0x62 |
| X | Execute | 0x58, 0x78 |
| P | Precursor | 0x50, 0x70 |
| D | Detach | 0x44, 0x64 |

#### Classification method

1. For each protein, scan all token byte sequences for the presence of any opcode byte value at any position within the token.
2. Apply a priority cascade: B → X → P → D. The first opcode detected determines the protein’s classification.
3. This mutual exclusivity ensures each protein receives a single, unambiguous opcode assignment.

Cross-species conservation of functional department composition was assessed across nine species spanning bacteria, archaea, and eukaryotes. Pairwise vocabulary similarity (Kendall’s τ over 22 shared functional departments) was correlated with evolutionary divergence time; results are reported in Results.

#### Functional Vocabulary (Computational Vocabulary)

The vocabulary was derived through systematic enrichment analysis of token-function associations across the human proteome.

#### Discovery pipeline

1. Token aggregation: All tokens from 62,796 human proteins were pooled, retaining the association between each token and its source protein(s).
2. Functional annotation mapping: Each source protein was mapped to its functional department annotations. Department labels were derived from UniProt Gene Ontology annotations (biological process, molecular function, cellular component) grouped into 32 functional categories representing major cellular programs.
3. Enrichment testing: For each token, Fisher’s exact test assessed whether proteins carrying that token were enriched for specific functional departments relative to the proteome background. P-values were corrected using Benjamini-Hochberg FDR.
4. Functional assignment: Tokens with significant enrichment (FDR < 0.05) were assigned to their dominant functional department. Confidence scores reflect annotation source: UniProt-curated (1.0), heuristic (0.70), and algorithmic convergence (0.36).

#### Vocabulary statistics

**Table 11.**
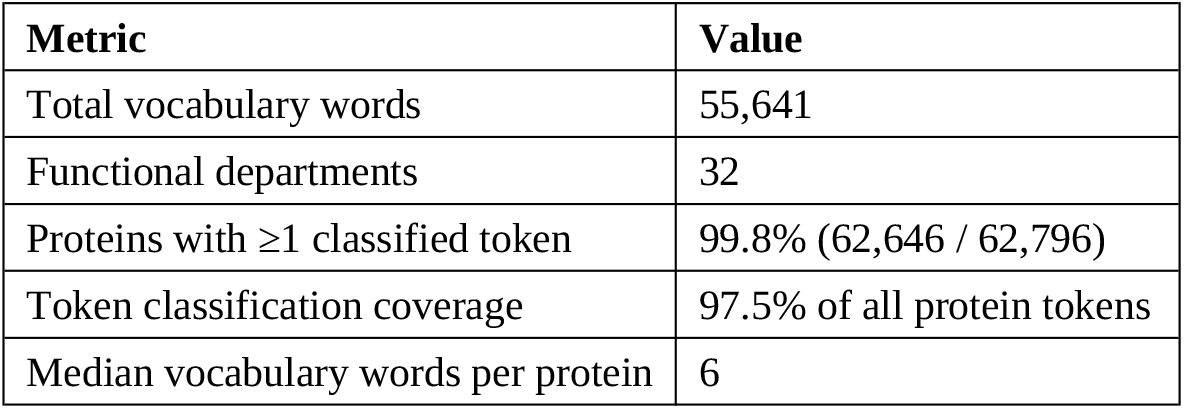
Vocabulary Statistics.

| Metric | Value |
| --- | --- |
| Total vocabulary words | 55,641 |
| Functional departments | 32 |
| Proteins with $\geq 1$ classified token | 99.8% (62,646 / 62,796) |
| Token classification coverage | 97.5% of all protein tokens |
| Median vocabulary words per protein | 6 |

The 32 departments span cellular functions including: Apoptosis, Autophagy, Cell adhesion, Cell cycle, Chromatin, Cytoskeleton, DNA repair, DNA replication, GTPase, Glycosylation, Immune response, Ion channel, Kinase, Lipid metabolism, Methylation, Mitochondrial, Nucleic acid binding, Nuclear transport, Olfactory, Phosphatase, Protein folding, Proteolysis, RNA processing, Receptor signaling, Signaling, Structural, Transcription, Translation, Transport, Ubiquitin, Uncharacterized, and Vesicle trafficking.

#### Cross-Validation Protocol

Vocabulary enrichment and function prediction were validated using a held-out protocol against UniProt-curated ground truth (confidence ≥ 0.5, retaining 28,075 proteins with high-quality functional annotations while excluding low-confidence algorithmic labels):

1. Split: The human proteome was randomly divided 50/50 into training and test sets (10 independent seeds, seeds 42–51).
2. Enrichment correlation: For each vocabulary word, enrichment was computed independently on training and test sets. Pearson correlation between train and test enrichments was calculated per seed.
3. Function prediction: For each protein, vocabulary words were aggregated by department, and the department with the highest cumulative confidence-weighted score was predicted. Predictions were evaluated against UniProt-curated ground truth using multi-label scoring (prediction correct if it matches any of the protein’s annotated departments).
4. Scaling analysis: Accuracy was stratified by vocabulary word count per protein to assess the relationship between token coverage and prediction performance.
5. Confidence calibration: The margin between the top-1 and top-2 prediction scores was assessed as a self-calibrating confidence metric.
6. Attractor analysis: Progressive removal of dominant prediction-target departments was performed to assess whether vocabulary discrimination improves when over-broad labels are excluded.

#### Chromosomal Convergence Validation

To validate that the protein-derived vocabulary reflects intrinsic genomic structure rather than an artifact of the back-translation pathway, the vocabulary was cross-referenced against patterns discovered independently from chromosomal DNA.

#### Method

1. All 24 nuclear chromosomes plus the mitochondrial genome were encoded via the direct DNA pathway (Steps 1–4).
2. Pattern discovery (Step 5) was applied, generating 27,677,556 genome patterns stored in the convergence_tier2 table.
3. Vocabulary intersection was computed by exact hash matching between protein vocabulary words and chromosome patterns.

Convergence rate increases with pattern length. This independent convergence validates that the vocabulary is encoded in the genome itself, not introduced by the protein encoding pathway.

### Step 7 – Program Discovery and Compositional Structure

The vocabulary-annotated patterns from Step 6 are assembled into genome programs through boundary detection and compositional analysis. This step identifies discrete functional units in both chromosomal and protein-derived sequences.

#### Genome Program Discovery

Programs are identified by detecting boundaries in the vocabulary signal - regions where functional annotation density drops sharply relative to adjacent positions.

#### Discovery method

1. Sliding window analysis: A fixed-width window (default: 50 bytes) traverses the encoded chromosome sequence. At each position, vocabulary hits are counted by matching window contents against the Computational Vocabulary vocabulary.
2. Boundary detection: Program boundaries are placed where the vocabulary density drops below a threshold relative to the local maximum (default: boundary_drop > 2.0×).
3. Entry point assignment: Each program is assigned an entry point - the first vocabulary pattern within its boundaries - recorded as chromosome:hex_position (e.g., chrM:0x5B2B).
4. Function sequence extraction: The ordered sequence of vocabulary functions within each program is recorded (e.g., Unclassified|Cytoskeleton|DNA repair).

#### Primitive Identification

Programs recurring across multiple chromosomes with identical function sequences are designated as primitives which are defined as fundamental operations conserved genome-wide.

#### Primitive criteria

1. Identical function sequence (ordered list of vocabulary departments)
2. Recurrence across ≥2 chromosomes
3. Minimum 3 distinct functions in the sequence

The 116 primitives represent the genome’s core instruction set - recurring functional patterns that appear across most or all chromosomes.

#### Compositional Nesting

Programs exhibit hierarchical structure: larger programs contain smaller programs as subsequences, forming a nesting hierarchy.

#### Nesting analysis

1. For each program pair, test whether the shorter program’s function sequence appears as a contiguous subsequence within the longer program.
2. Record nesting relationships with depth (number of enclosing layers).
3. Classify programs as Primitive (no nested subprograms) or Composite (contains one or more nested primitives).

Note: Current nesting analysis captures single-depth containment relationships. The 2,795 relationships represent direct parent-child nesting; multi-level nesting has not been assessed.

#### Execution Trace (Dispatch Graph)

Programs are connected through a dispatch graph recording which programs invoke or reference other programs. The dispatch graph is traced from the mitochondrial genome’s entry points, which serve as the kernel of the execution architecture.

#### Dispatch construction

1. For each mitochondrial entry point, identify all vocabulary patterns across all chromosomes that match the entry point’s patterns.
2. Record directed edges from source position to target position, annotated with the pattern function and hop depth.
3. Trace execution paths to specified hop depths (default: 2 hops).

#### Protein Program Synthesis

For the protein-derived pathway, each protein’s vocabulary words are assembled into a program representation:

1. Token aggregation: All vocabulary words from the protein’s token sequence are collected with their functional assignments.
2. Function sequence: The ordered list of vocabulary functions forms the protein’s program signature.
3. Complexity classification: Proteins are classified by vocabulary word count:
  a. Minimal (1–5 words)
  b. Simple (6–15 words)
  c. Moderate (16–50 words)
  d. Complex (51+ words)
4. Opcode annotation: The protein’s B/X/P/D opcode (from Step 6) is attached as execution metadata.

The resulting protein programs can be cross-referenced against genome programs via shared vocabulary patterns, enabling protein-to-genome functional mapping.

### Step 8 - Kernel Construction: Assembly of the Genomic Operating Kernel

The programs, primitives, and dispatch graph produced by Step 7 are assembled into a computational kernel, deemed the Operational Bioinformatics System (OBS) which is implemented as a multi-layer execution engine. The kernel provides the infrastructure for loading, validating, and dispatching the synthesized programs.

#### Architecture

The kernel comprises three layers:

1. Kernel core (obs/kernel/, 1,159 lines): Boot loader, instruction decoder, process table, signal dispatcher, and program executor. This is the runtime that loads genome data and executes programs.
2. OBS analysis modules (obs/, 2,918 lines): Higher-level analysis built on the kernel - cascade tracing, knockout simulation, gene resolution, and therapeutic intervention routing.
3. API layer (server/, 3,504 lines): TypeScript Express server providing the web-accessible interface, including real-time encoding pipeline execution, chromosome analysis, protein analysis, and VM simulation endpoints.

Total system: 7,581 lines (Python kernel + TypeScript API).

#### Boot Sequence and POST

The kernel employs a formal boot sequence modeled on hardware power-on self-test (POST) conventions. The boot loader is species-agnostic and discovers genome architecture from data rather than relying on hardcoded parameters.

Boot phases:

1. Phase 1 - POST (Power-On Self Test):
  a. Enumerate chromosomes from the genome data directory
  b. Load primitives (instruction set)
  c. Load programs (process images)
  d. Load nesting relationships
  e. Discover entry points from execution trace data
  f. Discover the kernel chromosome from entry point addresses
  g. Classify chromosome roles (KERNEL / RELAY / EFFECTOR) from cross-chromosome edge ratios using gap analysis
    i. Validate minimum requirements: instruction set size > 0, process count > 0, entry points > 0, chromosomes > 0
  h. Report boot OK or boot FAILED - no partial boots permitted
2. Phase 2 - Load Instruction Set:
  a. Initialize the instruction decoder with the primitives as opcodes
  b. Each opcode carries: rank, function sequence, classification, gene name, recurrence count, chromosome count
3. Phase 3 - Build Process Table:
  a. Create processes from programs, each assigned to a memory segment (chromosome) with protection level based on discovered role
  b. Protection levels: KERNEL_RO (read-only, kernel chromosome), RELAY_RW, RELAY_EFFECTOR_RW, EFFECTOR_RW
  c. Process states: LOADED (initial), ACTIVE (selected for execution), SIGNAL_ACTIVATED (reached by dispatch signal)
4. Phase 4 - Signal Dispatch:
  a. Load the dispatch graph
  b. Route signals from kernel entry points through discovered RELAY chromosomes to target processes
  c. Signal propagation model: strength = source_strength × exp(−λ × hop_depth) where λ is the decay constant
5. Phase 5 - Execute:
  a. Resolve program nesting grammar from the hierarchy data
  b. Decode instructions via the primitive instruction set
  c. Produce S-expression execution trees showing composite/primitive/opcode structure

#### Process Table and Memory Model

Each chromosome is treated as a memory segment with a protection level derived from its discovered role:

**Table 12.**
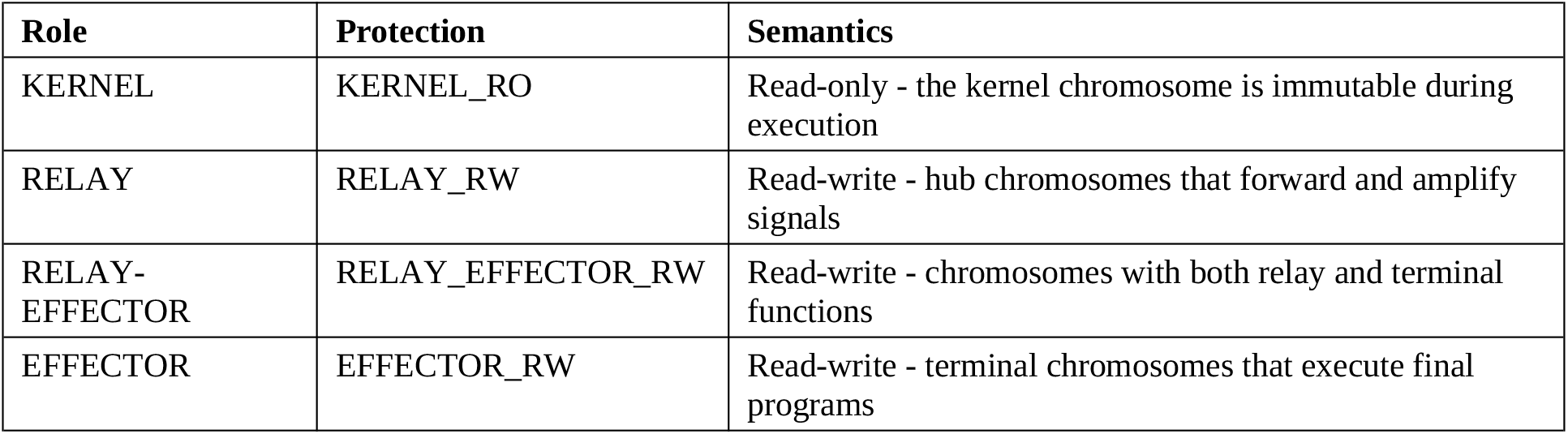
Chromosome Role Protection Levels.

The kernel chromosome has no processes in the process table because it is the kernel itself, not a process that runs on the kernel.

#### Execution Model

The kernel supports three levels of execution:

1. Level 1 (Structural): Resolves which primitives compose a program, which subroutines nest within it, and which processes are structurally reachable. Produces S-expression execution trees.
2. Level 2 (Weighted): Propagates signal strength through the dispatch graph with amplification and exponential decay. Produces per-process signal strength values.
3. Level 3 (Thermodynamic): Reserved - parameter slots exist in the codebase for thermodynamic modeling, but this level is not yet implemented.

#### Species Agnosticism

The kernel makes no assumption about which chromosome is the kernel, which is the relay hub, or how many chromosomes exist. All roles are discovered from data through gap analysis of cross-chromosome edge ratios. This design means the same kernel code can boot on any species’ genome data, provided the encoding pipeline (Steps 1–7) has been run on that species’ reference assembly. The only assumption is that one or more chromosomes serve as signal origins (entry points) and then from this point, the kernel discovers which ones from the dispatch graph.

#### Reproducibility

The kernel loads data from seven canonical CSV export files in the exports/ directory:

- programs_annotated.csv - genome programs
- primitive_annotations_complete.csv - primitives (instruction set)
- genome_subroutines_ranked.csv - subroutine definitions
- genome_nesting_hierarchy.csv - nesting relationships
- execution_trace_summary.csv - entry point edge counts
- execution_trace_hop1.csv - hop-1 dispatch edges
- execution_trace_graph_edges.csv.gz - hop-2 dispatch edges (compressed)

The kernel is stateless and deterministic. It loads data on startup and performs read-only analysis and underlying data is able to be modified by kernel operation. Re-running the boot sequence on the same export files produces identical output.

#### To boot the kernel from a terminal

1. python3 -m obs.kernel boot
2. Boot results for the human genome (chromosome role discovery, entry point statistics, relay hub identification, and signal dispatch architecture) are reported in Results.

#### Statistical Analysis

Five null model tests and one progressive peel analysis were conducted to validate the computational vocabulary produced by the encoding pipeline. Each test targeted a distinct structural claim: (i) that byte-stream patterns depend on amino acid ordering, not merely composition; (ii) that vocabulary-to-function mappings produce non-random protein-level convergence; (iii) that genome programs exhibit chromosome-specific concentration; (iv) that the inter-chromosomal dispatch network contains statistically significant hub structure; and (v) that vocabulary composition tracks phylogenetic distance. All tests used a fixed random seed (42) for reproducibility and 1,000 permutations unless otherwise noted. Empirical p-values were computed as (r + 1)/(n + 1), where r is the count of null statistics equal to or more extreme than the observed value and n is the number of permutations. Z-scores were computed as (observed − null mean)/null standard deviation. Multiple testing correction used Benjamini– Hochberg FDR < 0.05 where applicable. Effect sizes (Cohen’s d, τ^2^, odds ratios) are reported alongside p-values per current reporting guidelines.

#### Encoding null model

This test assessed whether the encoding pipeline produces byte and token distributions distinguishable from those generated by randomly permuted amino acid sequences. One thousand proteins were sampled from the human proteome (n = 78,007 proteins with sequences ≥50 amino acids). For each protein, the amino acid sequence was shuffled using Fisher–Yates randomization, preserving length and composition while destroying positional ordering. Both real and shuffled sequences were encoded through the deterministic pipeline: amino acid → codon (one fixed codon per amino acid) → DNA → 2-bit binary → hexadecimal bytes → tokens (contiguous 2–5 byte patterns occurring ≥2 times, top 100 retained per sequence). Three comparisons were performed: (i) byte frequency distributions compared using the two-sample Kolmogorov–Smirnov test and chi-square goodness-of-fit; (ii) vocabulary hit rates (tokens matching Computational Vocabulary words) compared using Welch’s t-test; (iii) per-protein Shannon entropy of byte distributions compared using Welch’s t-test.

#### Vocabulary convergence null model

This test evaluated whether functional labels assigned to vocabulary tokens produce meaningful protein-level convergence. For each of 93,296 proteins with Computational Vocabulary-matched tokens, a protein-by-function count matrix was constructed using SciPy sparse matrices (COO format). Functional overlap was defined as the mean HHI across proteins: for each protein, the fraction of token occurrences assigned to each functional label was squared and summed, yielding the probability that two randomly chosen tokens from the same protein share the same label. The null model consisted of 1,000 permutations in which primary_function labels were randomly shuffled across all 55,641 Computational Vocabulary token hashes, breaking the real token-to-function mapping while preserving the marginal distribution of labels and the protein-token structure. For each permutation, the protein-by-function matrix was reconstructed and mean overlap recomputed.

#### Primitive recurrence permutation test

This test assessed whether genome program motifs exhibit chromosome-specific concentration exceeding random expectation. Three statistics were computed: (i) per-chromosome maximum recurrence (highest count of any single function sequence within any chromosome); (ii) multi-chromosome spread (count of unique function sequences appearing on ≥3 chromosomes); (iii) chromosome concentration (mean HHI of chromosome distribution for each function sequence occurring ≥2 times). The null model preserved per-chromosome program counts while randomly reassigning function sequences. In each of 1,000 permutations, the pool of all function sequences was shuffled and redistributed to chromosomes in proportion to observed per-chromosome counts. P-values for maximum recurrence were one-sided (upper tail); p-values for spread and concentration were two-sided. Spearman correlation between primitive recurrence and chromosome count was computed across 116 annotated primitives.

#### Dispatch hub null model

This test evaluated whether the cross-chromosome dispatch network exhibits statistically significant hub structure. Three metrics were computed: (i) Gini coefficient of per-chromosome outbound edge weight sums; (ii) outbound-to-inbound ratio for chrM (mitochondrial chromosome); (iii) outbound-to-inbound ratio for chr19. The null model employed degree-preserving edge-swap randomization. In each of 1,000 permutations, 5,000 pairwise edge swaps were performed: two edges were selected at random and their source or target endpoints exchanged (swaps creating self-loops were rejected). This procedure approximately preserves each node’s total degree while randomizing topology. P-values were one-sided (upper tail), testing whether observed metrics exceeded the null distribution.

#### Cross-species conservation test

This test assessed whether opcode frequency distributions track evolutionary divergence. For nine species spanning ~3.5 billion years (*H. sapiens, M. musculus, D. rerio, C. elegans, D. melanogaster, S. cerevisiae, A. thaliana, E. coli, H. salinarum*), relative frequency of each classified opcode was computed from per-species vocabulary files. For all 36 pairwise comparisons (21 computable after excluding pairs with insufficient shared vocabulary), Kendall’s τ-b was computed on shared opcode frequency vectors. The primary statistic was the second-order correlation: Kendall’s τ between pairwise vocabulary similarity and established divergence times. Statistical significance was assessed analytically (scipy.stats.kendalltau) and by permutation (1,000 iterations shuffling pairwise τ values against fixed divergence times). Spearman’s ρ was computed as a complementary measure.

#### Progressive peel analysis

To assess robustness to dominant functional categories, three tests (vocabulary convergence, primitive recurrence, and cross-species conservation) were re-executed under progressive department exclusion. Three layers were defined: L0 (all 27 departments), L1 (Mitochondrial excluded), L2 (Mitochondrial and Transcription excluded). At each layer, Computational Vocabulary words belonging to excluded departments were removed prior to analysis. For vocabulary convergence, this reduced the dictionary from 55,641 to 48,567 (L1) and 37,789 (L2) words. For primitive recurrence, programs containing excluded departments in their function sequence were removed, reducing the corpus from 4,936 to 2,909 at L2. For cross-species conservation, vocabulary words with excluded labels were removed from each species’ frequency distribution. The encoding null model and dispatch hub null model were not included as their data sources are not indexed by functional department.

#### Functional enrichment analysis

FDR correction. For each of four opcodes (B/X/P/D), the proportion of proteins carrying that opcode and annotated to each Gene Ontology term was compared against proteome-wide background frequencies. Analyses were performed independently for H. sapiens, *S. cerevisiae*, and *E. coli*. Significant enrichments (FDR < 0.05) are reported with odds ratios and 95% confidence intervals. The B/X/P/D classification is a descriptive labeling scheme derived from byte-value ranges (see Supplementary Methods for full details); it has not been independently validated by a dedicated null model and its primary role is to provide coarse execution categories for downstream program annotation.

#### Vocabulary function prediction cross-validation

Function prediction accuracy was assessed using 10-fold stratified cross-validation with independent random seeds (42–51). The human proteome was split 50/50 into training and test sets per fold. Vocabulary-to-function mappings were learned on training proteins and evaluated on held-out test proteins. Multi-label accuracy was computed as the proportion of test proteins for which the top-predicted function matched any curated annotation. Only proteins with high-confidence UniProt annotations (confidence ≥ 0.5) were included, yielding 28,075 evaluation proteins. Pearson correlation between training and test enrichment values was computed per vocabulary word to assess generalization.

DepMap essentiality validation. To test whether vocabulary-derived functional assignments predict gene essentiality independently, mean Chronos scores from the DepMap 25Q3 CRISPR loss-of-function screen (1,186 cell lines, 18,435 genes) were obtained for each gene. Genes were matched to vocabulary department assignments by gene symbol, yielding 10,819 overlapping genes across 22 departments (each with 10 or more genes). Three statistics were computed: (i) eta-squared, measuring the fraction of Chronos score variance explained by department assignment (sum of squared between-group deviations divided by total sum of squares); (ii) Cohen’s d, comparing mean Chronos scores of genes in the five most essential departments (Translation, RNA processing, Cell cycle, Nucleic acid binding, DNA repair; n = 2,009) against all remaining departments (n = 8,810); (iii) Mann-Whitney U test for the same two-group comparison. DepMap data were not used at any stage of vocabulary construction, token assignment, or department classification.

## Supporting information

Supplementary Note 1: Worked Examples

## Acknowledgements

The author thanks M. Birlew for discussions on computational architecture and statistical methodology, and M. Jacobs (Metropolitan State University of Denver) for guidance on thermodynamic principles and scientific publishing. No external funding was received for this work.

## Author Contributions

J.L. conceived the study, designed the encoding pipeline, performed all analyses, wrote the kernel implementation, and wrote the manuscript.

## Data Availability

The human reference proteome (UP000005640) is available from UniProt (https://www.uniprot.org/proteomes/UP000005640). The human genome assembly (T2T-CHM13v2.0, GCA_009914755.4) is available from NCBI GenBank. Cross-species proteomes are available from UniProt under accession numbers listed in Methods. The computational vocabulary, genome programs, execution trace, and all validation datasets generated during this study are available at https://github.com/Omnis-Architecture-Co/genome-kernel-supplementary upon publication. Source data for all figures are provided with the paper.

## Code Availability

The OBS kernel and the complete encoding pipeline are available at https://github.com/Omnis-Architecture-Co/genome-kernel-supplementary upon publication. The pipeline is deterministic: re-execution on identical input produces byte-identical output. MD5 checksums for all input files are provided in Supplementary Information.

## References

1. Tanenbaum, A. S. & Bos, H. Modern Operating Systems 5th edn (Pearson, 2022).

2. Avsec, Ž. et al. Effective gene expression prediction from sequence by integrating long-range interactions. Nat. Methods 18, 1196–1203 (2021).

3. Avsec, Ž. et al. Advancing regulatory variant effect prediction with AlphaGenome. Nature 649, 1206–1218 (2026).

4. Nguyen, E. et al. Genome modelling and design across all domains of life with Evo 2. Nature (2026).

5. Almassalha, L. M. et al. Geometrically encoded positioning of introns, intergenic segments, and exons in the human genome. Adv. Sci. 13, e09964 (2025).

6. Yan, K.-K. et al. Comparing genomes to computer operating systems in terms of the topology and evolution of their regulatory control networks. Proc. Natl Acad. Sci. USA 107, 9186–9191 (2010).

7. Nurk, S. et al. The complete sequence of a human genome. Science 376, 44–53 (2022).

8. The UniProt Consortium. UniProt: the Universal Protein Knowledgebase in 2025. Nucleic Acids Res. 53, D199–D207 (2025).

9. Sayers, E. W. et al. GenBank 2025 update. Nucleic Acids Res. 53, D26–D31 (2025).

10. Ashburner, M. et al. Gene Ontology: tool for the unification of biology. Nat. Genet. 25, 25–29 (2000).

11. The Gene Ontology Consortium. The Gene Ontology knowledgebase in 2023. Genetics 224, iyad031 (2023).

12. Cunningham, F. et al. Ensembl 2024. Nucleic Acids Res. 52, D866–D872 (2024).

13. Martin, F. J. et al. Ensembl 2023. Nucleic Acids Res. 51, D933–D941 (2023).

14. Szklarczyk, D. et al. The STRING database in 2023: protein-protein association networks and functional enrichment analyses for any observed and predicted proteome. Nucleic Acids Res. 51, D7–D17 (2023).

15. Benjamini, Y. & Hochberg, Y. Controlling the false discovery rate: a practical and powerful approach to multiple testing. J. R. Stat. Soc. B 57, 289–300 (1995).

16. Virtanen, P. et al. SciPy 1.0: fundamental algorithms for scientific computing in Python. Nat. Methods 17, 261–272 (2020).

17. Harris, C. R. et al. Array programming with NumPy. Nature 585, 357–362 (2020).

18. Crick, F. H. C. Codon–anticodon pairing: the wobble hypothesis. J. Mol. Biol. 19, 548–555 (1966).

19. Church, G. M., Gao, Y. & Kosuri, S. Next-generation digital information storage in DNA. Science 337, 1628 (2012).

20. Dempster, J. M. et al. Chronos: a cell population dynamics model of CRISPR experiments that improves inference of gene fitness effects. Genome Biol. 22, 343 (2021).

21. Nielsen, A. A. K. et al. Genetic circuit design automation. Science 352, aac7341 (2016).

