## Supplementary Information for "A deterministic computational kernel encoded in the human genome"

Jasmine Levy

#### Contents

Supplementary Table 1. Standard genetic code with 6-bit binary encoding
Supplementary Table 2. Functional department classification scheme
Supplementary Table 3. Input file specifications and checksums
Supplementary Table 4. Cross-species vocabulary checksums
Supplementary Table 5. Reproducibility parameters
Supplementary Note 1. Worked example: nucleolin (protein-derived pathway)
Supplementary Note 2. Worked example: chromosome 11 INS locus (DNA-native pathway)
Supplementary Data. Source data files (deposited at https://github.com/Omnis-Architecture-Co/genome-kernel-supplementary)

#### Supplementary Table 1

**Standard genetic code with 6-bit binary encoding.** Each amino acid is mapped to its first-listed RNA codon per the standard genetic code, then to DNA via U→T substitution. The resulting 6-bit binary value (2 bits per nucleotide: A=00, T=01, G=10, C=11) determines the byte stream used in all downstream analyses. Synonymous codons sharing first-position nucleotides produce identical 6-bit values (e.g., both AAA and AAG encode Lys as 001000), ensuring robustness to codon degeneracy. Stop codons are excluded from protein-derived encodings but retained in chromosomal pathway encodings. The complete 64-codon table with binary values is provided in the source data (TableS5_codon_encoding.csv).

| Codon | Amino Acid | 1-letter | 6-bit Value | Binary | Hex |
|---|---|---|---|---|---|
| ATG | Met (start) | M | 10 | 001010 | 0x0A |
| GCT | Ala | A | 16 | 010000 | 0x10 |
| AAG | Lys | K | 8 | 001000 | 0x08 |
| GAG | Glu | E | 3 | 000011 | 0x03 |
| CTT | Leu | L | 9 | 001001 | 0x09 |
| TAA | Stop | * | — | — | — |

*Representative rows shown. Complete 64-codon table available in source data.*

#### Supplementary Table 2

**Functional department classification scheme.** Each vocabulary word was assigned to one of 27 functional departments (plus Unclassified) through Fisher's exact test enrichment analysis against Gene Ontology annotations with Benjamini–Hochberg FDR correction. Departments are listed in descending order by word count.

| Department | Words | % of Vocabulary | Mean Enrichment | Example Words |
|---|---|---|---|---|
| Transcription | 525 | 27.2% | 6.34 | 0xC4DA |
| Unclassified | 438 | 22.7% | — | 0x75D5 |
| Chromatin | 241 | 12.5% | 5.63 | 0x8DF4 |
| Structural | 149 | 7.7% | 17.98 | 0x882A |
| Cytoskeleton | 127 | 6.6% | 6.16 | 0x8693 |
| RNA processing | 112 | 5.8% | 12.91 | 0xE5DE |
| Cell adhesion | 62 | 3.2% | 7.98 | 0x0610 |
| Cell cycle | 58 | 3.0% | 4.62 | 0x0620 |
| DNA repair | 42 | 2.2% | 4.01 | 0x096A |
| Nucleic acid binding | 34 | 1.8% | 3.71 | 0x0BD0 |
| Methylation | 29 | 1.5% | 11.50 | 0x0B2C |
| Phosphatase | 27 | 1.4% | 6.86 | 0x4144 |
| Kinase | 21 | 1.1% | 4.61 | 0x0A9B |
| Transport | 14 | 0.7% | 3.98 | 0x0AD1 |
| Ion channel | 11 | 0.6% | 4.73 | 0x35D1 |
| GTPase | 10 | 0.5% | 6.23 | 0xE410 |
| Translation | 9 | 0.5% | 12.16 | 0x082B |
| Ubiquitin | 8 | 0.4% | 9.25 | 0x1041 |
| Protein folding | 4 | 0.2% | 12.97 | 0x8421 |
| Proteolysis | 4 | 0.2% | 6.65 | 0x5A11 |
| Apoptosis | 3 | 0.2% | 6.97 | 0xE72C |
| Signaling | 3 | 0.2% | 3.83 | 0x5B90 |
| Immune | 1 | 0.1% | 5.80 | 0x5A159D |

#### Supplementary Table 3

**Input file specifications and checksums.** MD5 and SHA256 checksums enable verification that identical inputs produce identical outputs. All files are available in the source data repository.

| File | Rows | MD5 Checksum |
|---|---|---|
| vocabulary.csv (human) | 1,932 | e57e9cd0e0a0722f210e40e5abb2eb1d |
| programs_annotated.csv | 4,936 | 23285bb0d53c05b4f1b30b8f7bbef86d |
| primitive_annotations_complete.csv | 116 | a73f48eab42ce10ad280ca13e3745f8e |
| execution_trace_summary.csv | 126 | c3d960c5f65c60bd5f598391f6f9a7d6 |
| execution_trace_hop1.csv | 43,554 | ce6e8c090541c9a5d1b4f65f339f82f3 |
| chromosome_roles.csv | 24 | e8a52bbdde6fc49c5ab1cd00568b9fa0 |
| species_registry.json | 6 species | 1ce1fa359af8acd2848cf804fbfb1ff4 |
| genome_primitives.csv | 116 | ff30248dce5d096d05b543b9e6e32e5e |

#### Supplementary Table 4

**Cross-species vocabulary checksums.** Per-species vocabulary files used in cross-species conservation analysis (VAL-XSP-001). All species names are italicised in the main text per taxonomic convention.

| Species | Words | MD5 Checksum |
|---|---|---|
| H. sapiens | 1,932 | e57e9cd0e0a0722f210e40e5abb2eb1d |
| M. musculus | 1,117 | fc4829219ad8f7093edbde009b710556 |
| D. rerio | 2,403 | 2c8e764e1a23e956dc79cef0c2591209 |
| D. melanogaster | 937 | 23250b6f95b5cabc2c557502c137d029 |
| S. cerevisiae | 123 | ef858e4e5b27b515d17fc71b4f8d6f6b |
| E. coli | 7 | c6f907f07fe93e1b5e4421d6548e6933 |

### Supplementary Table 5

**Reproducibility parameters.** All random number generators were seeded with value 42. Each validation test produces a JSON results file with full provenance (file hashes, row counts, parameters), a Markdown summary, and a PNG visualisation with annotated statistics.

| Parameter | Value |
|---|---|
| Pipeline | 6-bit codon encoding (OIP v2) |
| Random seed | 42 |
| Permutations (all null models) | 1,000 |
| Cross-validation folds | 10 (seeds 42–51) |
| Kernel boot command | python3 -m obs.kernel boot |
| Validation suite command | python3 -m validation.run_all --seed 42 |
| Sliding window (programs) | 50 bytes |
| Boundary drop threshold | 2.0× |
| Primitive min chromosomes | ≥2 |
| Primitive min distinct functions | ≥3 |
| Edge swaps per permutation (NET-001) | 5,000 |
| Token pattern length | 2–5 bytes (protein); 2–100 bytes (chromosome) |
| Min token frequency | ≥2 |

#### Supplementary Note 1

**Worked example: nucleolin (protein-derived pathway).** The human nucleolin protein (UniProt P19338, gene NCL, 710 amino acids) was carried through all pipeline stages as a reproducibility verification example. Nucleolin is the major nucleolar protein of growing eukaryotic cells, found associated with intranucleolar chromatin and pre-ribosomal particles. Its 710-residue length produces 532 complete bytes, sufficient for statistically meaningful byte distributions. The complete step-by-step encoding (amino acid → RNA codon → DNA → binary → bytes → hex → tokens) is documented in Methods Steps 1–4 of the main manuscript, with the first six residues (M-V-K-L-A-K) traced through each stage.

Key properties of the nucleolin encoding: 532 complete bytes (4-bit remainder discarded); byte range distribution 10.3% Control (0x00–0x1F), 43.0% Standard (0x20–0x7F), 46.6% Extended (0x80–0xFF). The first complete byte (0x1A, decimal 26) falls in the Control range, reflecting the universal ATG start codon signature shared by all protein-derived encodings.

#### Supplementary Note 2

**Worked example: chromosome 11 INS locus (DNA-native pathway).** A 1,000-nucleotide segment of chromosome 11 (GRCh38 coordinates 11p15.5) spanning the INS gene and its flanking regulatory regions was carried through all pipeline stages as the chromosomal-substrate reproducibility example. Unlike the protein pathway, this example begins with raw genomic DNA (including introns, untranslated regions, and intergenic sequence) and applies nucleotide-to-binary encoding directly without the amino acid → codon intermediary.

Key properties of the INS locus encoding: 250 complete bytes (no remainder); byte range distribution 9.2% Control, 38.0% Standard, 52.8% Extended. The higher proportion of Extended bytes relative to the protein-derived example (52.8% vs 46.6%) reflects the greater nucleotide diversity of non-coding genomic sequence. Despite distributional differences, 77 of the 127 unique byte values (60.6%) present in the nucleolin encoding also appear in the chromosomal encoding, indicating substantial vocabulary overlap between the two independent substrates.

#### Supplementary Data

The following source data files are deposited at https://github.com/Omnis-Architecture-Co/genome-kernel-supplementary and are available upon publication:

**Vocabulary and classification:** Complete 1,932-word human vocabulary dictionary with functional annotations, enrichment values, p-values, and carrier gene lists (vocabulary_human_1932words.csv). Six cross-species vocabulary files for mouse (1,117 words), zebrafish (2,403), fly (937), yeast (123), and E. coli (7). Gene-to-department mapping (gene_departments.csv).

**Genome programs and primitives:** All 4,936 annotated genome programs with chromosome, position, function sequence, entry point, recurrence, and gene annotations (programs_annotated_4936.csv). All 116 primitive annotations with BLAST results and genomic positions (primitive_annotations_116.csv).

**Dispatch architecture:** Execution trace summary (7 entry points with dispatch edge counts), 43,554 hop-1 dispatch edges, 552 cross-chromosome connection pairs with edge counts (11.4M total edges), and chromosome role classifications.

**Validation outputs:** JSON results files for all validation tests (encoding null model, vocabulary convergence, primitive recurrence, dispatch hub, cross-species conservation, progressive peel analysis) with complete provenance metadata including file hashes, row counts, and parameters.
