## Supplementary Note 1: Worked Examples for "A deterministic computational kernel encoded in the human genome"

### Encoding Pipeline Worked Examples

This note traces two biological sequences through the complete 8-step encoding pipeline described in Methods. Example A follows nucleolin (UniProt P19338), a 710-amino-acid nucleolar phosphoprotein, through the protein-derived pathway. Example B follows a 1,000-nucleotide segment of the chromosome 11 INS locus (T2T-CHM13v2.0, 11p15.5) through the DNA-native pathway. Both examples use real pipeline outputs at every stage.

*Encoding rule (both pathways): each DNA nucleotide maps to a 2-bit value (A = 00, T = 01, G = 10, C = 11). For proteins, each amino acid is first reverse-translated to its OMNIS first-listed DNA codon (Table S5), producing a 6-bit code per residue (3 nucleotides × 2 bits). The resulting bit stream is then packed into 8-bit bytes.*

### Example A: Nucleolin (Protein-Derived Pathway)

Nucleolin (gene NCL, UniProt P19338) is a 710-amino-acid nucleolar phosphoprotein involved in ribosome biogenesis, chromatin remodeling, and transcription regulation. Its N-terminal domain contains alternating acidic and basic stretches; its C-terminal domain contains four RNA recognition motifs (RRMs) and a glycine/arginine-rich (GAR) domain.

#### Step 1 — Amino Acid to Codon to Binary Encoding

Each amino acid is reverse-translated to its OMNIS first-listed DNA codon (Table S5), then each nucleotide of the codon is converted to its 2-bit binary value (A = 00, T = 01, G = 10, C = 11), yielding 6 bits per residue. The first six residues of nucleolin (MVKLAK) are encoded as follows:

| Position | Residue | Amino Acid | DNA Codon (Table S5) | Binary (6-bit) |
|---|---|---|---|---|
| 1 | M | Methionine | ATG | 000110 |
| 2 | V | Valine | GTT | 100101 |
| 3 | K | Lysine | AAG | 000010 |
| 4 | L | Leucine | CTT | 110101 |
| 5 | A | Alanine | GCT | 101101 |
| 6 | K | Lysine | AAG | 000010 |

The complete 710-residue sequence produces 4,260 bits of binary data (710 residues × 3 nucleotides × 2 bits).

#### Step 2 — Binary-to-Byte Conversion

The 4,260-bit stream is partitioned into 8-bit bytes, yielding 532 complete bytes with a 4-bit remainder that is discarded (not zero-padded), preventing spurious terminal patterns. Because each amino acid contributes 6 bits, the byte boundaries do not align with residue boundaries—each byte contains parts of adjacent codons.

| Byte | Hex | Decimal | Bit Pattern | Source Bits |
|---|---|---|---|---|
| 1 | 0x1A | 26 | 00011010 | from MV codons |
| 2 | 0x50 | 80 | 01010000 | from VK codons |
| 3 | 0xB5 | 181 | 10110101 | from KL codons |
| 4 | 0xB4 | 180 | 10110100 | from AK codons |

### Step 3 — Byte Range Classification

Each byte is classified into one of three ranges based on its value:

| Range | Hex Values | Count | Percentage |
|---|---|---|---|
| Control | 0x00–0x1F | 55 | 10.3% |
| Standard | 0x20–0x7F | 226 | 42.5% |
| Extended | 0x80–0xFF | 251 | 47.2% |
| Total | — | 532 | 100.0% |

The distribution (10% Control / 42% Standard / 47% Extended) reflects the codon-based encoding: G-initial codons (binary prefix 10) and C-initial codons (prefix 11) push bytes toward the Extended range, while A-initial codons (prefix 00) produce Control-range bytes.

### Step 4 — Hexadecimal Representation

Each byte is expressed as a two-character hexadecimal value. Where the byte falls in the printable ASCII range (0x20–0x7F), the corresponding character is shown for reference:

| Byte | Hex | Decimal | ASCII | Range |
|---|---|---|---|---|
| 1 | 0x1A | 26 | · | Control |
| 2 | 0x50 | 80 | P | Standard |
| 3 | 0xB5 | 181 | · | Extended |
| 4 | 0xB4 | 180 | · | Extended |
| 5 | 0x2B | 43 | + | Standard |
| 6 | 0x69 | 105 | i | Standard |
| 7 | 0x08 | 8 | · | Control |
| 8 | 0x1C | 28 | · | Control |

*Complete hex stream (first 40 characters):*
*1A50B5B42B69081CA987D0821ADF7DF428A58A28…*

### Step 5 — Token Discovery

The tokenizer scans the byte stream with a greedy longest-match algorithm, matching multi-byte patterns against the 1,932-word vocabulary (word lengths range from 2 to 5 bytes). For nucleolin's 532-byte stream, the scanner identifies 93 vocabulary hits across 76 unique words.

| Position | Word (hex) | Length (bytes) | Occurrences in corpus | Department |
|---|---|---|---|---|
| 3 | 0xB42B | 2 | 80 | Unclassified |
| 9 | 0x87D0 | 2 | 101 | Cytoskeleton |
| 13 | 0xDF7D | 2 | 4223 | Chromatin |
| 15 | 0xF428 | 2 | 160 | Cytoskeleton |
| 18 | 0x8A28 | 2 | 2841 | Chromatin |
| 20 | 0x5D8A18 | 3 | 21 | Unclassified |
| 26 | 0xA18A28 | 3 | 41 | Chromatin |
| 30 | 0x75DA | 2 | 346 | Chromatin |
| 32 | 0x628A | 2 | 2934 | Chromatin |

| | | | | |
|---|---|---|---|---|
| 37 | 0x2082 | 2 | 1456 | Chromatin |
| 39 | 0xA420 | 2 | 155 | Unclassified |
| 42 | 0xB6D3 | 2 | 170 | Chromatin |
| 45 | 0xB420 | 2 | 181 | DNA repair |
| 51 | 0x3420 | 2 | 211 | DNA repair |
| 58 | 0xD082 | 2 | 1193 | Chromatin |
| … | (61 more) | … | … | … |

### Step 6 — Vocabulary Classification

Each matched vocabulary word carries a functional department assignment derived from GO-term enrichment analysis of its carrier proteins. Nucleolin's 93 vocabulary hits map to the following department distribution:

| Department | Hits | Percentage |
|---|---|---|
| Chromatin | 29 | 31.2% |
| Unclassified | 26 | 28.0% |
| RNA processing | 9 | 9.7% |
| DNA repair | 8 | 8.6% |
| Cytoskeleton | 5 | 5.4% |
| Cell adhesion | 4 | 4.3% |
| Transcription | 3 | 3.2% |
| Nuc acid bind | 3 | 3.2% |
| Structural | 2 | 2.2% |
| Kinase | 1 | 1.1% |
| Translation | 1 | 1.1% |
| Cell cycle | 1 | 1.1% |
| Methylation | 1 | 1.1% |

The dominant classified department is Chromatin (29 of 93 hits, 31%), consistent with nucleolin's known role in chromatin remodeling and histone chaperone activity. The Unclassified category (26 hits) represents general-purpose vocabulary words that are functionally promiscuous across multiple departments. The protein receives a vocabulary density of 17.48% (hits per byte).

### Step 7 — Program Assembly

In the protein-derived pathway, each protein's vocabulary words are assembled into a program representation. Nucleolin's program has the following properties:

| Property | Value |
|---|---|
| Gene | NCL (Nucleolin) |
| UniProt | P19338 |
| Vocabulary words | 93 |
| Unique words | 76 |
| Distinct departments | 13 |
| Complexity tier | Complex |
| Dominant classified dept | Chromatin |
| Function sequence (first 80 chars) | Unclassified\|Cytoskeleton\|Chromatin\|<br>Cytoskeleton\|Chromatin\|Unclassified\|Chromati… |

The function sequence records the ordered list of departments encountered along the protein's byte stream. This sequence serves as the protein's program signature for cross-referencing against genome programs.

#### Step 8 — Kernel Integration

At the kernel level, nucleolin's protein program is indexed by its UniProt accession and gene name. The kernel's process table records:

| Property | Value |
|---|---|
| Protein PID | P19338 (NCL) |
| Chromosome | chr2 (NCL gene locus: 2q37.1) |
| Memory segment | RELAY_EFFECTOR_RW |
| Vocabulary density | 17.48% |
| Program type | Protein-derived |
| Dispatch connections | Via shared vocabulary words with genome programs |

Nucleolin's vocabulary words can be cross-referenced against genome programs carrying the same multi-byte patterns, establishing protein-to-genome dispatch connections. For example, vocabulary word 0xDF7D (Chromatin) appears in 4223 proteins genome-wide, linking nucleolin to the broader Chromatin regulatory network.

### Example B: Chromosome 11 INS Locus (DNA-Native Pathway)

The INS locus (T2T-CHM13v2.0 chr11, cytogenetic band 11p15.5) encodes the insulin gene. This 1,000-nucleotide segment spanning the INS gene and its flanking regulatory regions is traced through the DNA-native encoding pathway, which differs from the protein pathway in Step 1: nucleotides are encoded directly (2 bits each) rather than through reverse translation of amino acids (6 bits each via codon lookup).

#### Step 1 — Nucleotide-to-Binary Encoding

Each nucleotide is assigned a 2-bit binary value using the same mapping as the codon pipeline: A = 00, T = 01, G = 10, C = 11. The first 12 nucleotides of the INS locus region are:

| Position | Nucleotide | Binary |
|---|---|---|
| 1 | G | 10 |
| 2 | C | 11 |
| 3 | A | 00 |
| 4 | T | 01 |
| 5 | C | 11 |
| 6 | T | 01 |
| 7 | G | 10 |
| 8 | A | 00 |
| 9 | T | 01 |
| 10 | T | 01 |
| 11 | C | 11 |
| 12 | C | 11 |

The 1,000-nucleotide sequence produces 2,000 bits of binary data.

#### Step 2 — Binary-to-Byte Conversion

With 2 bits per nucleotide, exactly four nucleotides fill one byte (4 × 2 = 8 bits). The 1,000 nucleotides yield exactly 250 bytes with no remainder.

| Byte | Hex | Decimal | Source Nucleotides | Bit Pattern |
|---|---|---|---|---|
| 1 | 0xB1 | 177 | GCAT | 10110001 |
| 2 | 0xD8 | 216 | CTGA | 11011000 |
| 3 | 0x5F | 95 | TTCC | 01011111 |
| 4 | 0x6C | 108 | TGCA | 01101100 |

### Step 3 — Byte Range Classification

| Range | Hex Values | Count | Percentage |
|---|---|---|---|
| Control | 0x00–0x1F | 20 | 8.0% |
| Standard | 0x20–0x7F | 98 | 39.2% |
| Extended | 0x80–0xFF | 132 | 52.8% |
| Total | — | 250 | 100.0% |

The DNA-native pathway produces a markedly different byte distribution than the protein pathway: 53% Extended versus nucleolin's 47%. This reflects the 2-bit encoding scheme, where GC-rich regions (G = 10, C = 11) produce high byte values.

### Step 4 — Hexadecimal Representation

| Byte | Hex | Decimal | ASCII | Range |
|---|---|---|---|---|
| 1 | 0xB1 | 177 | · | Extended |
| 2 | 0xD8 | 216 | · | Extended |
| 3 | 0x5F | 95 | _ | Standard |
| 4 | 0x6C | 108 | l | Standard |
| 5 | 0xC8 | 200 | · | Extended |
| 6 | 0x67 | 103 | g | Standard |
| 7 | 0x6A | 106 | j | Standard |
| 8 | 0xCF | 207 | · | Extended |

*Complete hex stream (first 40 characters):*
*B1D85F6CC8676ACF2B5BDE7577E37FC3DEC765A2…*

### Step 5 — Token Discovery

Scanning the 250-byte INS stream against the 1,932-word vocabulary (using greedy longest-match) yields 5 hit(s) across 5 unique word(s). The substantially lower hit rate compared to nucleolin (5/250 = 2.00% vs 93/532 = 17.48%) reflects a fundamental difference between the two pathways: the vocabulary was extracted from protein-derived encodings, so DNA-native sequences produce sparser vocabulary matches. Genome programs are identified by regions of concentrated vocabulary hits across entire chromosomes, not uniform coverage of short windows.

| Position | Word (hex) | Length (bytes) | Department | Occurrences |
|---|---|---|---|---|
| 36 | 0xB7DA | 2 | Unclassified | 213 |
| 153 | 0x28A2 | 2 | Chromatin | 3738 |
| 166 | 0x2882 | 2 | Chromatin | 1552 |
| 237 | 0x688D | 2 | Cell cycle | 260 |
| 243 | 0xC967 | 2 | Chromatin | 105 |

### Step 6 — Vocabulary Classification

The 5 vocabulary hit(s) in the INS region map to the following department(s):

| Department | Hits |
|---|---|
| Chromatin | 3 |
| Unclassified | 1 |
| Cell cycle | 1 |

### Step 7 — Program Assembly

Genome programs are identified by scanning complete chromosomes for regions where vocabulary density exceeds the boundary detection threshold. Within 500 kb of the INS locus, the pipeline identifies 4 genome programs:

| PID | Range | Length (bytes) | Function Sequence | Vocab Hits | Entry Point |
|---|---|---|---|---|---|
| chr11:1842232 | 1842232–1842313 | 82 | Chromatin\|Cell adhesion\|Transcription | 21 | chrM:0x71C7 |
| chr11:2126445 | 2126445–2126590 | 146 | Unclassified\|Chromatin\|Cytoskeleton\|Transc… | 135 | chrM:0x71C7 |
| chr11:2186858 | 2186858–2186891 | 34 | Unclassified\|Chromatin\|Cytoskeleton\|Chroma… | 51 | chrM:0x5B2B |
| chr11:2494696 | 2494696–2494790 | 95 | Cell cycle\|Chromatin\|Unclassified\|Kinase\|C… | 55 | chrM:0x5B2B |

Of these, 4 program(s) match known primitives (recurring cross-chromosomal patterns). The closest program to the INS gene (chr11:2126445) carries 6 distinct functional departments and is classified as REGULATORY.

### Step 8 — Kernel Integration

At the kernel level, each genome program near the INS locus receives a process ID (PID) based on its chromosomal position. These programs are registered in the kernel’s process table:

| PID | Role | Memory Segment | Entry Point | Dominant Function |
|---|---|---|---|---|
| chr11:1842232 | RELAY-EFFECTOR | RELAY_EFFECTOR_RW | chrM:0x71C7 | NAMED_GENE |
| chr11:2126445 | RELAY-EFFECTOR | RELAY_EFFECTOR_RW | chrM:0x71C7 | REGULATORY |
| chr11:2186858 | RELAY-EFFECTOR | RELAY_EFFECTOR_RW | chrM:0x5B2B | REGULATORY |
| chr11:2494696 | RELAY-EFFECTOR | RELAY_EFFECTOR_RW | chrM:0x5B2B | NAMED_GENE |

All four programs trace their dispatch origin to chrM entry points (0x71C7 and 0x5B2B), confirming that the INS region is reachable from the kernel’s boot sequence. Chromosome 11 is classified as RELAY-EFFECTOR with read-write memory protection.

### Substrate Convergence: Protein and DNA Pathways

A central feature of the V2 encoding pipeline is that both protein sequences and genomic DNA are encoded into the same byte space, enabling direct comparison. The protein pathway reverse-translates amino acids to DNA codons before applying the same nucleotide-to-binary mapping (A = 00, T = 01, G = 10, C = 11) used by the DNA pathway. The following analysis compares the encoding outputs of nucleolin (protein-derived) and the INS locus (DNA-native).

#### Byte-Level Comparison

| **Property** | **Nucleolin (protein)** | **INS Locus (DNA)** | **Shared** |
|---|---|---|---|
| Input length | 710 amino acids | 1000 nucleotides | — |
| Encoding path | AA → codon → DNA → binary | DNA → binary | — |
| Bits per input unit | 6 (3 nt × 2 bit) | 2 | — |
| Encoded bytes | 532 | 250 | — |
| Unique byte values | 125 | 144 | 75 (39% of union) |
| Control range | 10.3% | 8.0% | — |
| Standard range | 42.5% | 39.2% | — |
| Extended range | 47.2% | 52.8% | — |
| Vocabulary hits | 93 | 5 | — |
| Hit rate | 17.48% | 2.00% | — |

#### Vocabulary-Level Comparison

Nucleolin’s encoding contains 76 unique vocabulary words; the INS locus contains 5. They share no vocabulary words, which reflects the different byte distributions produced by the two encoding schemes. The protein-derived pathway (6-bit codon-based encoding) produces byte values distributed across all three ranges, while the DNA-native pathway (2-bit nucleotide encoding) concentrates in the Extended range due to the high information content per nucleotide pair.

#### Pathway Relationship

The two pathways intersect at the vocabulary level: both produce byte streams that are scanned against the same 1,932-word dictionary. When a vocabulary word appears in both a protein’s encoding and a genomic region’s encoding, it establishes a cross-substrate link. For the INS locus specifically, the genome programs identified within 500 kb of the gene carry function sequences (Chromatin, Transcription, Cytoskeleton) that overlap with departments found in nucleolin’s encoding, illustrating how the shared vocabulary enables cross-substrate functional annotation.

This convergence is not guaranteed by the method — it is an empirical result. The vocabulary was extracted from protein data; that it produces biologically meaningful hits in genomic DNA is a validation of the encoding’s ability to bridge the two substrates.
